# Rescue of blood coagulation Factor VIII exon-16 mis-splicing by antisense oligonucleotides

**DOI:** 10.1101/2023.03.31.535160

**Authors:** Victor Tse, Guillermo Chacaltana, Martin Gutierrez, Nicholas M. Forino, Arcelia G. Jimenez, Hanzhang Tao, Phong H. Do, Catherine Oh, Priyanka Chary, Isabel Quesada, Antonia Hamrick, Sophie Lee, Michael D. Stone, Jeremy R. Sanford

## Abstract

The human *Factor VIII* (*F8*) protein is essential for the blood coagulation cascade and specific *F8* mutations cause the rare bleeding disorder Hemophilia A (HA). Here, we investigated the impact of HA-causing single-nucleotide mutations on *F8* pre-mRNA splicing. We found that 14/97 (∼14.4%) coding sequence mutations tested in our study induced exon skipping. Splicing patterns of 4/11 (∼36.4%) *F8* exons tested were especially sensitive to the presence of common disease-causing mutations. RNA-chemical probing analyses revealed a three-way junction structure at the 3′ end of intron 15 (TWJ-3-15). TWJ-3-15 sequesters the polypyrimidine tract, a key determinant of 3′ splice site strength. Using exon-16 of the *F8* gene as a model, we designed specific antisense oligonucleotides (ASOs) that target TWJ-3-15 and identified three that promote the splicing of *F8* exon-16. Interaction of TWJ-3-15 with ASOs increases accessibility of the polypyrimidine tract and inhibits the binding of hnRNPA1-dependent splicing silencing factors. Moreover, ASOs targeting TWJ-3-15 rescue diverse splicing-sensitive HA-causing mutations, most of which are distal to the 3’ splice site being impacted. The TWJ-3-15 structure and its effect on mRNA splicing provide a model for HA etiology in patients harboring specific *F8* mutations and provide a framework for precision RNA-based HA therapies.

## INTRODUCTION

Noncoding sequences (introns) interrupt protein coding information (exons) in most human genes. Conserved sequences known as splice sites (ss) demarcate exon-intron boundaries (1). Messenger RNA (mRNA) biogenesis requires intron removal from precursor transcripts and exon ligation (2, 3). The spliceosome, a multi-megadalton ribonucleoprotein (RNP) complex assembles *de novo* on every intron to catalyze splicing reactions. This process involves the stepwise assembly of five uracil-rich small nuclear ribonucleoprotein particles (U snRNPs) and hundreds of accessory RNA-binding proteins (4–5). Exon definition is an initial spliceosome assembly step where splice site recognition occurs (6). In this early spliceosome complex, the U1 snRNP recognizes the 5′ss while the U2 snRNP auxiliary factor (U2AF) binds the 3′ss and polypyrimidine tract (7–10). This initial step is highly regulated in cells by the presence or absence of sequences that can function as exonic splicing enhancers or silencers (11–16).

Aberrant splicing contributes to the etiology of many inherited diseases (17). Disease-causing mutations often antagonize splicing by the direct ablation of splice site sequences and exonic splicing enhancors (ESEs), or the creation of exonic splicing silencers (ESSs). These lesions cripple the integrity of the gene by resulting in the production of a faulty message (18–20). Antisense oligonucleotides (ASOs) are a promising therapeutic modality for rescuing pathogenic aberrant splicing patterns as their direct base-pairing abilities make them highly customizable and specific to targets. Although challenges such as toxicity, delivery and stability represent barriers to the clinical translation of ASOs (21), solutions to these challenges exist, as exemplified by the recent FDA approval of multiple ASO drugs (22). Examples of these include the splice-modulating drugs Nusinersen and Milasen, where Nusinersin was the first FDA approved cure for spinal muscular atrophy (23–25) and Milasen is a patient-specific ASO for treatment of Batten’s disease (26).

Our previous work implicated thousands of disease-causing mutations in aberrant splicing (18–17). Among these, the *F8* gene had the highest frequency of mutations predicted to affect splicing regulatory sequences (Sterne-Weiler 2014). The *F8* gene encodes a protease required for initiation and activation of the coagulation cascade. *F8* deficiency causes Hemophilia A (HA), a deadly, X-linked recessive bleeding disorder. In some cases, aberrant splicing of the *F8* pre-mRNA contributes to HA etiology (27–30). In this study, we tested the impact of 97 HA-causing mutations on *F8* pre-mRNA splicing and discovered a structured RNA element in the intron upstream of exon-16 that attenuates the 3′ss. Interestingly, ASOs targeting and mitigating this structure rescue diverse splicing-sensitive mutations in *F8* exon-16. Together, these data reveal an unexpected RNA structure-function relationship modulating exon identity and provide a framework for the development of novel ASO therapeutics to treat HA.

## MATERIAL AND METHODS

### *F8* Splicing Reporters

The sequences of wild-type (WT) *F8* exons and 100-250 nucleotides flanking each exon were amplified from human genomic DNA (Promega) using WT PCR primers shown in Supplemental Table 1. Following gel purification, PCR products were ligated into pACT7_SC14 (*HBB* minigene reporter as previously described (31) using homology-based cloning technology (In-Fusion HD Cloning kit, Takara Bio). Following sequence verification, each plasmid was then used as a template for site-directed mutagenesis via overlap-extension PCR using mutagenesis primers shown in Supplemental Table 1. Mutant (MT) splicing reporter constructs were then sequence-validated using Sanger sequencing to confirm successful cloning and identity of splicing reporters. The designation for each *F8* point mutation, and therefore each MT *F8* exon presented in this study, is based on the nucleotide being mutated (e.g., A>C), and its position within the sequence context tested (i.e., length of flanking introns included and size of exon tested).

### Cell-based *in vivo* Splicing Assays

HEK293T cells (ATCC) were cultured in 6-well tissue culture plates (CytoOne, USA Scientific) using Dulbecco’s Modified Eagle Medium (Gibco, supplemented with 10% FBS) at 37°C, 5% CO_2_. The cells were transiently transfected at ∼60-80% confluency with 2.5ug of each *F8* splicing reporter using Lipofectamine 2000 (Invitrogen). Total RNA was harvested from cells 24-hours post-transfection using the Direct-zol RNA Miniprep kits (Zymo Research). Each *in vivo* splicing assay was performed a minimum of three times.

### ASO Walk and Combinatorial ASO Experiments

2′-methoxyethyl (2’MOE) phosphorothioate substituted ASOs complementary to *F8* exon-16 and flanking introns were designed from the reverse complement of the *F8* sense sequence, creating non-overlapping 18-mers as shown in Supplemental Table 2. *F8* exon-16 ASOs were designed to contiguously tile across the exon and its flanking introns. ASOs were synthesized by Integrated DNA Technologies (IDT). Each ASO is designated by their complementary positions in the *F8* exon-16 reporter. HEK293T cells (ATCC) were cultured in 96-well tissue culture plates (Perkin Elmer) as described above. Cells were transiently transfected with 250 ng of WT or MT splicing reporter and 10 pmol of each ASO using Lipofectamine 2000 (Invitrogen). 24-hours post transfection, cells were harvested and prepared for total RNA purification using the Quick-DNA/RNA Viral MagBead kit from Zymo Research and an Agilent Bravo NGS A liquid handler(32). Each experiment type (e.g., ASO walk or combinatorial ASO assays) was performed a minimum of three times.

### hnRNPA1 overexpression and western blot analysis

HEK293T cells (ATCC) were cultured in 6-well tissue culture plates as described above. Cells were co-transfected with 1.25 ug of the WT splicing reporter, 1.25 ug of either an empty expression vector or a T7-tagged hnRNPA1 expression vector, and 50 pmol of ASO(s) as described above. Total RNA and protein were isolated 24-hours post transfection using a RSB lysis buffer (10mM Tris pH 7.0, 100mM NaCl, 5mM MgCl2, 0.5% NP40, 0.5% Triton X-100, and EDTA-free Protease Inhibitor Cocktail (Roche)). Following a 10-minute incubation on ice, the cell lysate was then centrifuged at 10,000 x g for 10 minutes at 4°C. The supernatant was then collected and aliquoted for two separate applications. The first aliquot, comprising ∼90% of the cell lysate, was prepared for total RNA purification using the Direct-zol RNA Miniprep kits from Zymo Research. The remaining ∼10% of the cell lysate was then homogenized into a denaturing buffer solution containing 4X NuPAGE™ LDS Sample Buffer in preparation for polyacrylamide gel electrophoresis (Invitrogen™ NuPAGE™, 4 to 12%, Bis-Tris, 1.0–1.5 mm, Mini Protein Gels), and subsequent Western blots. For western blots, protein samples were transferred to a Immobilon NC membrane (Millipore) using a Genie Blotter (Idea Scientific). Membranes were probed with anti-HSP90 (Santa Cruz Biotech) and anti-T7 (Novagen) monoclonal antibodies and visualized by HRP conjugated secondary antibodies and chemiluminescence (Pierce). These experiments were performed a minimum of three times.

### Two-step RT-qPCR and Analysis of Splicing Reporter Assays

A minimum of 500 ng of purified total RNA was used as input for all first-strand cDNA synthesis using Multiscribe Reverse Transcriptase (Applied Biosystems). The resulting cDNA was then used as a template for endpoint PCR amplification using specific primers that detect our mRNA splicing reporter isoforms. The forward primer of the pair contains a 5′ FAM modification. The resulting amplicons were then analyzed using agarose gel electrophoresis to empirically evaluate mRNA isoforms detected. The abundance of each 5′ FAM labeled mRNA isoform is quantified using capillary electrophoresis and fragment analysis (UC Berkeley, DNA Sequencing Center). For fragment analysis, each sample is suspended in a formamide solution that contains a proper size standard for sizing detected fragments (GeneScan 1200 Liz, Applied Biosystems). Analysis was performed in PeakScanner (Thermofisher).

### Calculating Splicing Efficiency using Percent-Spliced-In (PSI) Index Formula

Based on fragment analysis data collected, subsequent quantification of splicing efficiency is achieved by comparing relative fluorescence units (RFU) between 5′ FAM labeled reporter isoforms that include or exclude an exon of interest. The RFU detected for each reporter isoform is then plugged into the following formula to calculate the PSI index, which reflects the splicing efficiency of an exon in either the WT or MT context:

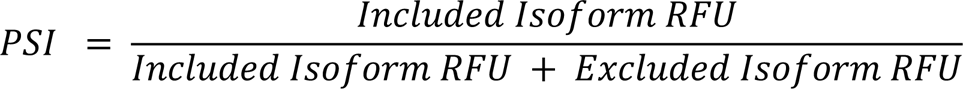

The mean PSI for a given reporter context is then calculated using all its respective replicates for a corresponding experiment. Statistical significance in the differences between the mean PSI of the control group(s) vs the experimental group(s) is determined using analysis of variance (ANOVA), and Dunett’s *post-hoc* test. All statistical tests for PSI analysis were done in GraphPad Prism 9. Values are determined to be statistically significant if calculated the *P*-value is below an alpha value of ≤ 0.05.

### *In vitro* Transcribed RNA for *F8* exon-16 WT and Its MTs

Templates for WT or MT *F8* exon-16 pre-mRNA sequences corresponding to the reporter plasmid inserts were synthesized by a primer assembly reaction designed using Primerize (33). RNA was purified by denaturing PAGE and eluted from gel slices overnight in 10 mM Tris pH 7.5, 480 mM sodium acetate, 1 mM EDTA, 0.1% SDS. Following ethanol precipitation transcripts were resuspended in ddH2O and quantified by UV spectrophotometry.

### *In vitro* SHAPE Probing of *F8* Targets

*F8* exon-16 *in vitro* transcribed pre-mRNA sequences were first denatured by incubating at 95°C for 3 minutes in 65 mM Na-HEPES (pH 8.0). The denatured RNA was then allowed to slowly cool to room temperature (RT) for 15 minutes, after which MgCl_2_ was supplemented to 1 mM for a total volume of 15 µL and incubated at RT for an additional 5 minutes. To chemically modify RNA, 2-aminopyridine-3-carboxylic acid imidazolide (2A3) was added to a final concentration of 100 mM and incubated for 2 minutes at 37°C (34, 35). The reaction was then quenched using dithiothreitol (DTT) to a final concentration of 500 mM at RT for 10 minutes. Reactions which substituted anhydrous dimethyl sulfoxide (DMSO) for 2A3 were used as negative controls. All modified RNAs, including negative controls, were then purified using RNA Clean & Concentrator-5 (Zymo Research).

### SHAPE-MaP Library Assembly and Sequencing

Modified RNAs were fragmented to a median size of 200 nucleotides by incubation at 94°C for 1 minute using NEBNext® Magnesium RNA Fragmentation Kit and then purified using NEB’s recommended ethanol precipitation protocol. Purified RNA was then prepared for reverse transcription, incubating the RNA with 1µL of 10mM dNTPs and 2µL of 20 µM random hexamers at 70°C for 5 minutes, followed by immediate transfer to ice. Reverse transcription reactions were then supplemented with 4µL of 5X RT buffer (250 mM Tris-HCl pH 8.3, 375 mM KCl), 2 µL of 0.1 M DTT, 1 µL of 120 mM MnCl2, 10 U of SUPERase RNase Inhibitor, and 200 U of Superscript II Reverse Transcriptase (SSII) [ThermoFisher Scientific, cat. 18064014] to a final volume of 20 µL. These reactions were then incubated at 25°C for 10 minutes to allow for partial primer extension, followed by incubation at 42°C for 3 hours to enable efficient extension. SSII was then heat-inactivated by incubation at 75°C for 20 minutes. Reverse transcription reactions were then supplemented with EDTA to a final concentration of 6 mM to chelate Mn^2+^ ions and incubated at RT for 5 minutes. MgCl_2_ was then added to a final concentration of 6mM for each reaction (34). Reverse transcription reactions were then used as input material for NEBNext® Ultra™ II DNA library Prep Kit for Illumina® (New England Biolabs, cat. E7645L), using NEBNext Multiplex Oligos for Illumina® (Unique Dual index UMI Adaptors DNA Set 1, cat. E7395). Subsequent reactions were performed following manufacturer instructions. All sequencing was performed on an Illumina iSeq100 instrument using a paired end 2×150 sequencing reagent cartridge and flow cell.

### SHAPE-MaP Data Analysis and RNA Structure Prediction

All the relevant data analysis steps were conducted using RNA Framework v2.6.9 (36). Reads produced from Illumina libraries were pre-processed and mapped using the rf-map module (parameters: -b2 -mp “--no-mixed –no-discordant” -bs) ensuring only paired-end mates with expected mate orientation were considered with Bowtie2. The mutational signal was obtained using the rf-count module (parameters: -m -pp -nd -ni) enabling mutation counts of reads produced from properly paired mates. Mutational signal was normalized relative to an unmodified control using parameters (-sm 3 -nm 1 - mu 0.05) and further normalized using the 2-8% normalization approach provided by RNA Framework (37). Normalized reactivities were then supplied to RNAstructure to generate data-driven predicted structure models (38).

### In silico analysis of protein-RNA interactions and splice site strength

MaxEntScan was used to analyze splice 5’ and 3’ site sequences of exons used in this study (39). To determine 3’ss strength, 23-mer sequences encompassing the last 20 intronic positions and first three exonic positions were selected for analysis. For 5’ss strength 9-mers consisting of the last three positions of each exon and the first six positions of the downstream intron were selected for analysis. The tool RBPmap (40) was used to identify putative protein-RNA interactions sites within *F8* pre-mRNAs. In this analysis, we used sequences corresponding to *F8* exon-16 and flanking introns from the splicing reporter. We included a high stringency constraint to match known RBP motifs within our input sequence, as well as a conservation filter to selectively identify motifs that best match sequences from the human and mouse genomes.

## RESULTS

### Multiple F8 exons exhibit aberrant splicing in the presence of HA-causing mutations

We previously described inherited disease-causing mutations with the potential to alter the landscape of ESEs or ESSs (18). Among all candidate genes, *F8* had the highest number of mutations per exon and total number of putative splicing-sensitive point mutations(17). In this study, we investigated the impact of a wide array of HA-causing mutations on the splicing of eleven *F8* exons (Fig. 1A). To determine whether these HA-causing mutations can induce aberrant splicing, we analyzed 97 distinct mutations by generating heterologous splicing reporters where the wild type (WT) or mutant sequences for each *F8* exon plus 100-250 bp of flanking intron sequence were cloned into the *beta globin (HBB*) minigene splicing reporter (Fig. 1B). Following transient transfections of each reporter into HEK293T cells, *HBB* reporter mRNA isoform levels are quantified by a two-step end-labeled RT-PCR assay and capillary electrophoresis. Out of the eleven *F8* exons tested, we found that HA-causing mutations in four exons (exon-7, exon-11, exon-16, and exon-18) caused exon skipping in the heterologous reporter context (Supplemental Fig. 1), indicating that the splicing fidelity of these exons may be particularly sensitive to mutations (Fig. 1C; Supplemental Fig 2). For example, Fig. 1C shows splicing assays for 16 HA-causing variants of exon-16. In comparison to the WT exon-16 splicing reporter which is efficiently spliced (lane 3), we found that six pathogenic mutations, namely G351A, C348T, C321A, C321T, C179T, and A333G significantly reduced exon-16 inclusion (Fig. 1C, compare lane 3 to lanes 5, 7, 10, 11, 14 and 18; Fig. 1D). Taken together, these results show that HA-causing mutations can alter *F8* pre-mRNA splicing efficiency and that exon-16 appears to be particularly sensitive to such mutations.

**Figure 1.**
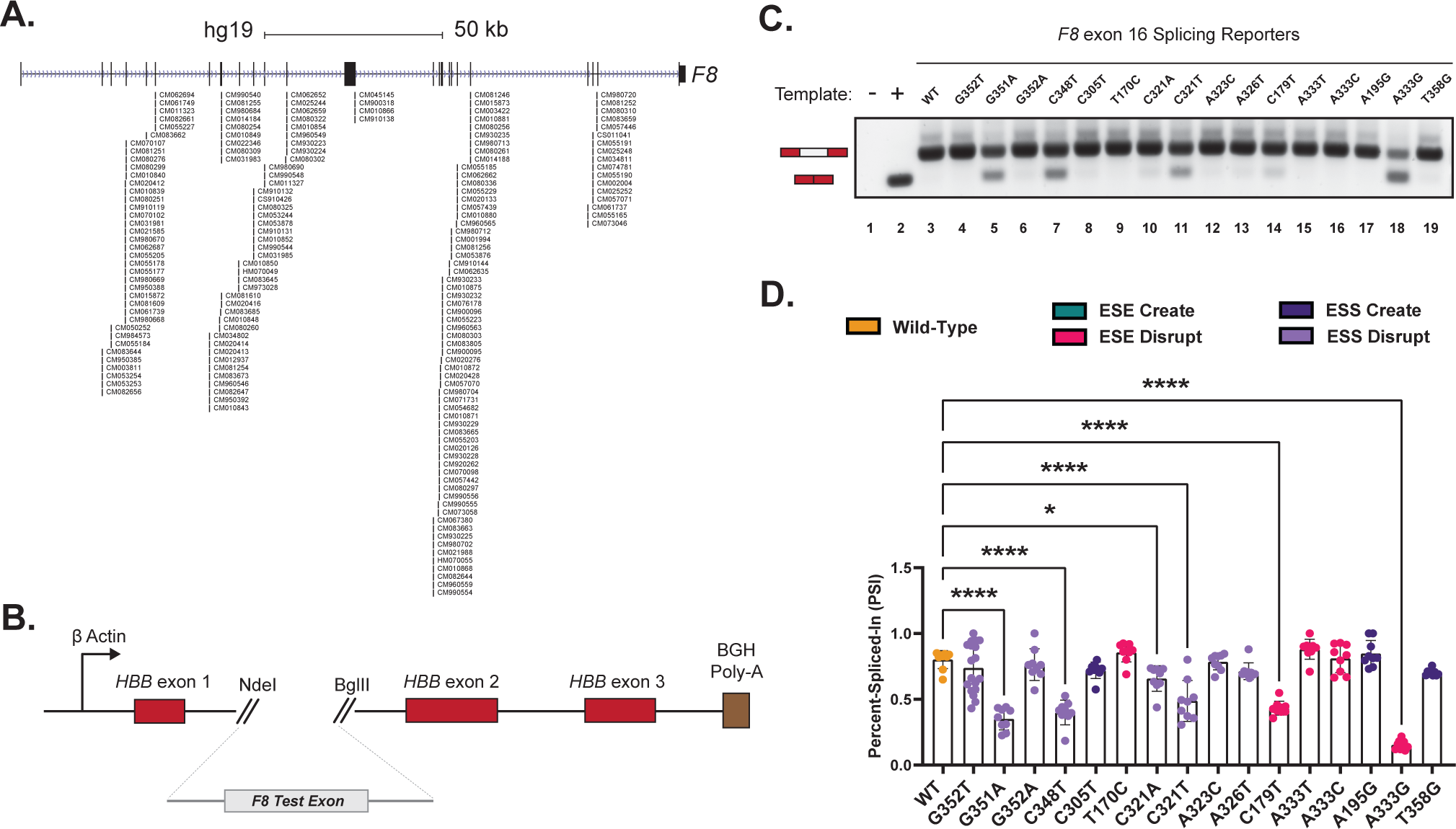
*In vivo* splicing reporter assays reveal a highly fragile exon susceptible to mutation-induced aberrant splicing. (**A**) *F8* gene structure schematic showing the positions and Human Gene Mutation Database (HGMD) IDs of 97 known HA-causing mutations tested in this study. The chevrons overlapping the introns (thin lines) point in a 5′ to 3′ orientation. Exons are designated by closed boxes. (**B**) Schematic of the heterologous splicing reporter used to assess the impacts of mutations on splicing. Each test exon and flanking intronic sequence of *F8* gene was cloned between exons 1 and 2 of the *HBB* minigene. (**C**) A representative agarose RNA gel showing the effects on splicing by various HA-causing mutations in a panel of *F8* exon-16 splicing reporters. Controls include a no template reaction (lane 1) and a positive control for exon skipping (lane 2). (**D**) Quantification of various HA-causing mutations on the splicing extent of *F8* exon-16. Percent-spliced-in (PSI) refers to the ratio of test exon skipped to test exon included in mRNA. Each mutation’s predicted impact on regulatory elements involved in splicing are annotated by color. Statistical significance between comparisons is denoted by asterisks that represent *P*-values with the following range of significance: **\*** = *P* ≤ 0.05, **\*\*** = *P* ≤ 0.01, **\*\*\*** = *P* ≤ 0.001, **\*\*\*\*** = *P* ≤ 0.0001.

### Modulation of F8 exon-16 splicing by antisense oligonucleotides (ASOs)

Mutations in *F8* exon-16 induce aberrant splicing. To determine if splicing of *F8* exon-16^A333G^ can be corrected by ASOs, we designed non-overlapping, phosphorothioate-substituted, 2′-methoxyethyl modified 18-mer ASOs that span exon-16 and its flanking introns in a splicing reporter context (Fig. 2A; Supplemental Table 2). In all ASO walk experiments, we include an ASO that has no complementarity to *F8* sequences, serving as our non-targeting (NT) control (Fig. 2B, lane 1), and an ASO “blocker” that specifically targets the 5′ss of exon-16 to directly inhibit its splicing (Fig. 2B, lane 2). We discovered multiple ASOs that enhanced splicing of exon-16^A333G^ (Fig. 2C). Co-transfection of ASO_1-18_, ASO_37-54_, ASO_55-72_, ASO_91-108_, or ASO_469-486_ with exon-16^A333G^ resulted in a statistically significant increase in exon-16 inclusion relative to its control with no ASOs co-transfected. Of note, these ASOs target the upstream and downstream intronic regions of exon-16^A333G^. ASO_1-18_, ASO_37-54_, ASO_55-72_, and ASO_91-108_ target regions upstream and adjacent to the 3′ss, rescuing splicing respectively by ∼1.5-fold (*P*-value = 0.0182), ∼2.0-fold (*P*-value <0.0001), ∼1.8-fold (*P*-value = <0.0001), and ∼1.9-fold (*P*-value = <0.0001). ASO_469-486_ targets a region downstream of the 5′ss, rescuing splicing by ∼1.7-fold (*P*-value = 0.0007). Taken together, our ASO walk indicates that targeting regions primarily upstream of *F8* exon-16 with ASOs may rescue splicing patterns of splicing-sensitive mutants by perturbing the influence of inhibitory elements found in the flanking introns.

**Figure 2.**
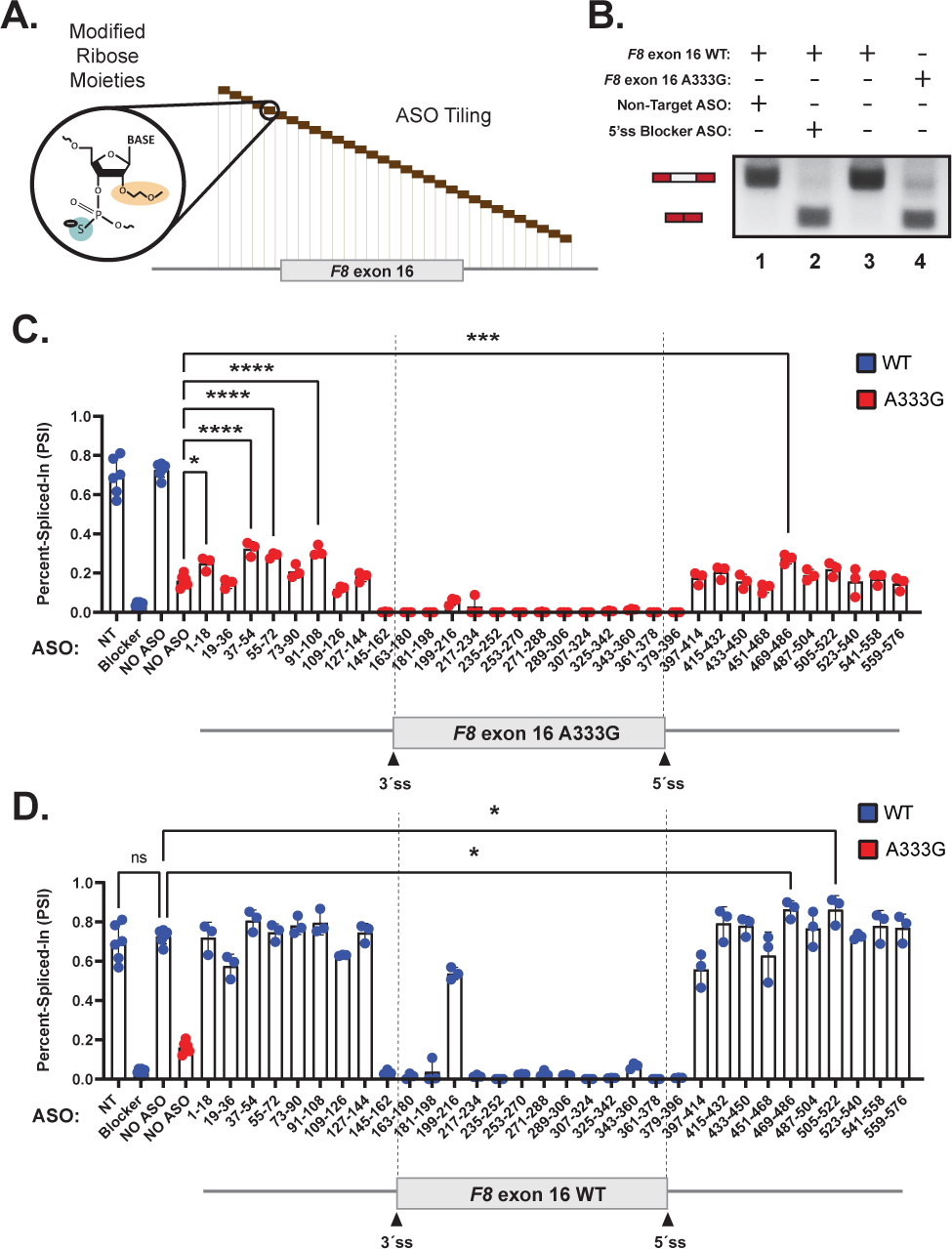
ASO walk reveals splice-modulating ASOs for *F8* exon-16 and a highly splicing-sensitive mutation in exon-16. (**A**) A mock schematic of an ASO walk. Each ASO used in our walks are 18 nucleotides in length and are designed using ribose sugars that are modified 2′-methoxyethyl group (2′-MOE, highlighted in light orange), and the phosphate backbone is modified to a phosphorothioate backbone (highlighted in light blue). Each 18-mer ASO is contiguous by design, tiling across exon-16 and its flanking introns with no overlaps between each ASO. (**B**) Proof-of-concept demonstrating how our ASOs and controls are expected to work in the ASO walk experiments. As shown in the annotative matrix above a representative agarose gel, the first two controls consist of our 5’ss blocker ASO (positive control) and our non-targeting ASO (negative control) being co-transfected with our WT exon-16 splicing reporter to demonstrate that our designed ASOs can modulate splicing. The last two controls consist of our WT exon-16 and exon-16^A333G^ mutant splicing reporters without ASOs co-transfected to illustrate the typical splicing ratios we may expect to see from their splicing. Expected mRNA isoforms including or excluding the test exon are also annotated to the left of the agarose gel (**C**) and (**D)** show our ASO walk data for the exon-16^A333G^ mutant and WT exon-16, respectively. The WT context is annotated by a blue color whereas the A333G mutant is annotated by a red color. ASO walk results for both (**C**) and (**D**) are quantified using the PSI ratio. Statistical significance between comparisons are denoted by asterisks that represent *P*-values with the following range of significance: ns = *P* > 0.05, **\*** = *P* ≤ 0.05, **\*\*** = *P* ≤ 0.01, **\*\*\*** = *P* ≤ 0.001, **\*\*\*\*** = *P* ≤ 0.0001. A schematic model of exon-16 and its flanking introns at the bottom of each plot to illustrate relative positions of ASOs.

As a control we repeated the ASO walk on the WT *F8* exon-16 reporter. As observed for the exon-16^A333G^ reporter, ASOs directly targeting the WT exon-16 reporter, except for ASO_199-216_, strongly inhibited its splicing (Fig. 2D). By contrast, individual ASOs targeting the flanking introns had little impact on exon-16 splicing relative to our controls. Two ASOs that target the intronic region downstream of the 5′ss, ASO_469-486_ and ASO_505-522_, are indicated to be statistically significant in enhancing splicing relative to the WT control with no ASO. ASO_469-486_ results in a 1.2-fold increase in splicing (*P*-value = 0.0106), whereas ASO_505-522_ also results in a 1.2-fold increase in splicing (*P*-value = 0.0114). ASOs that target the flanking intronic sequences upstream of the 3′ss appear to modulate splicing positively or negatively in a subtle, non-statistically significant manner.

### SHAPE-MaP-seq reveals a splicing-inhibitory RNA structure near the 3′ss of F8 exon-16

Disease-causing mutations have previously been reported to disrupt native RNA structure and as a consequence, alter their biological function (41). To determine how splicing-sensitive mutations influences RNA structures in *F8* exon-16, we performed selective 2′-hydroxyl acylation analyzed by primer extension and mutational profiling coupled to high-throughput sequencing (SHAPE-MaP-seq) on *in vitro* transcribed RNA. All *in vitro* RNA corresponding to WT exon-16 or its HA variants contain the same sequence context as our splicing reporters (34). Accessible nucleotides strongly react with 2-aminopyridine-3-carboxylic acid imidazolide (2A3) in SHAPE-MaP-seq assays (34). Nucleotides acylated by 2A3 can vary in their degree of modification, or reactivity, reflecting how accessible the nucleotides may be in the context of the global folding architecture of the RNA (34, 37, 42). We generated SHAPE reactivity profiles for WT exon-16 and the splicing-sensitive exon-16^A333G^ variant (Fig. 3A-B). We used arc diagrams to compare SHAPE-driven folding predictions of WT and exon-16^A333G^ transcripts. The exon-16^A333G^ variant induces subtle RNA structural changes, creating several long-range base-pairing interactions within the exon and between the flanking introns (Fig. 3C). Like exon-16^A333G^, other splicing-sensitive HA mutations in exon-16 exhibited modest structural changes compared to the WT transcript (Supplemental Fig. 3 and 4). Taken together, the SHAPE-MaP-Seq experiments suggest that splicing sensitive HA-causing mutations modestly impact the secondary structure of exon-16 compared to WT. Intriguingly, when analyzing both the WT and mutant SHAPE-driven structure predictions together, we discovered a three-way junction RNA structure in the intronic region upstream of *F8* exon-16 near the 3′ ss that is supported by the experimental reactivity patterns but not from in silico RNA folding alone (Fig. 3D-E and Supplemental Fig. 5). Our SHAPE-driven structural predictions suggest that an upstream region of *F8* intron-15 base-pairs with the branchpoint and polypyrimidine (poly-Y) tract, potentially occluding their accessibility to splicing factor 1 (SF1) and U2AF. Comparison of our SHAPE and ASO walk data revealed that ASOs which individually improved splicing of exon-16^A333G^ directly target this structure. These ASOs include ASO_55-72_ and ASO_91-108_, which had a statistically significant improvement on exon-16^A333G^ splicing, and ASO_73-90_ which also targets the structure but was unable to significantly rescue splicing of exon-16^A333G^. This 3’ss three-way junction structure, which we will refer to as TWJ-3-15 (**<u>T</u>**hree-**<u>W</u>**ay **<u>J</u>**unction at the **<u>3</u>**′ end of intron **<u>15</u>**), may be important for rescuing splicing-sensitive mutations in exon-16. Taken together, our data suggest that co-transfecting more than one ASO targeting TWJ-3-15 may further destabilize this structure to increase 3′ss accessibility and thereby restore proper splicing of exon-16.

**Figure 3.**
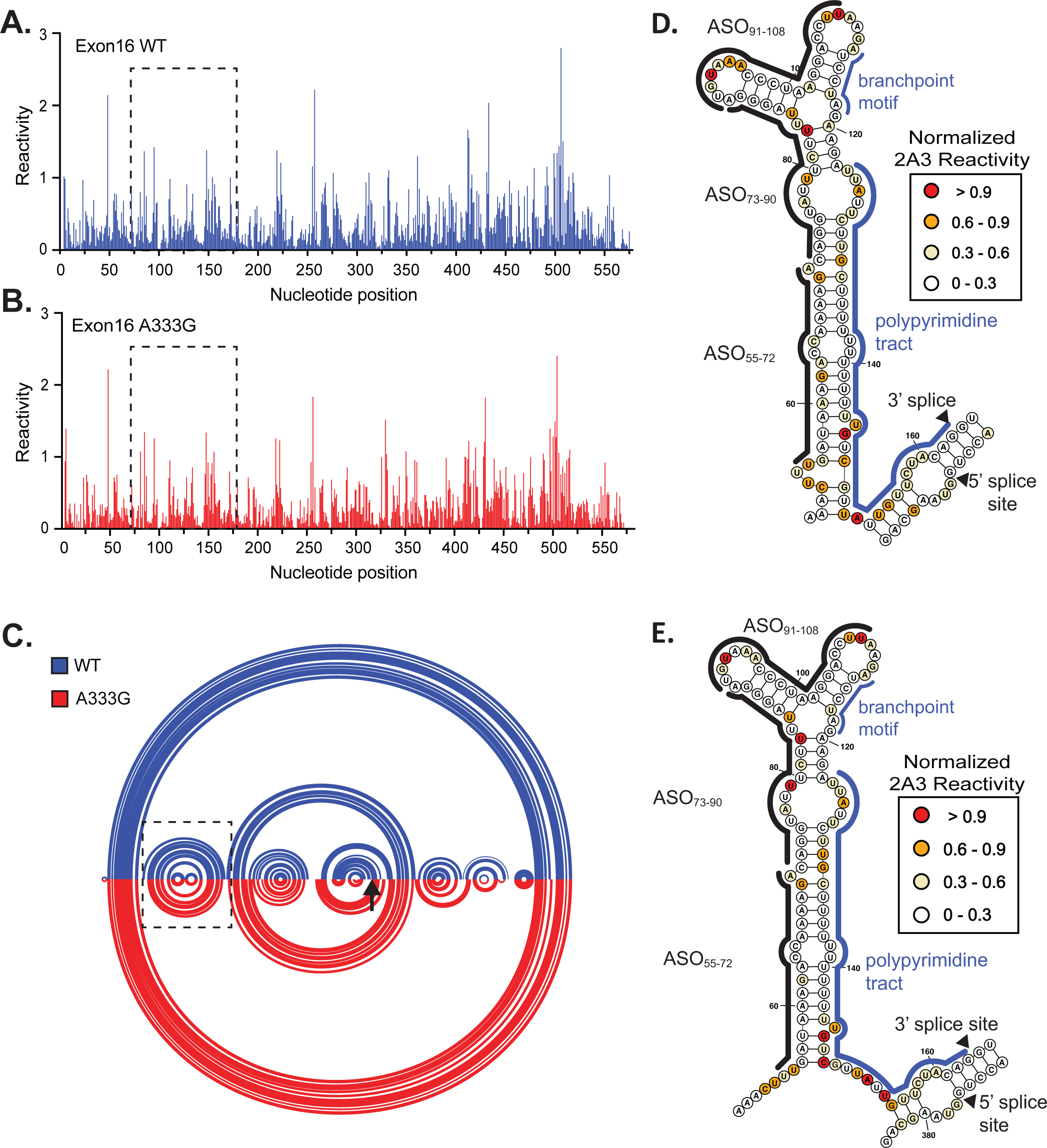
SHAPE probing identifies a native RNA structure (TWJ-3-15) that is uniquely positioned at the 3′ss of *F8* exon-16. (**A, B**) A normalized SHAPE reactivity plot for WT exon-16 (blue) and the exon-16^A333G^ mutant (red), respectively. **(C)** Intramolecular base pairing interactions, constrained by normalized SHAPE reactivity are represented by arcs joining different regions of the transcript. Arc diagrams for WT and A333G mutant transcript are depicted in blue and red, respectively. The broken box indicates the position of TWJ-3-15. (**D, E**) SHAPE-driven secondary structure prediction of TWJ-3-15 depicted in its two dimensional state for WT and A333G exon-16 transcripts, respectively. Core splicing signals, *cis*-regulatory elements, and ASOs are also annotated within the structure. All SHAPE probing data presented were generated *in vitro* using the SHAPE reagent 2A3, and all subsequent data analysis was performed in RNA Framework. The sequence is numbered according to the nucleotide position from the 5′ to 3′ orientation.

### Combinations of ASOs additively enhance F8 exon-16 splicing as seen in the exon-16^A333G^ variant by destabilizing TWJ-3-15

To test the hypothesis that a combination of ASOs can additively enhance splicing of *F8* exon-16^A333G^ compared to a single ASO when targeting TWJ-3-15, we performed additional *in vivo* splicing assays where exon-16^A333G^ is co-transfected with ASO_55-72_, ASO_73-90_ and ASO_91-108_. Remarkably, we discover that this trio ASO combination had a striking effect on exon-16^A333G^ splicing (Fig. 4A). Compared to the exon-16^A333G^ control without any ASOs co-transfected (Fig. 4A, lane 4), in addition to a duo combination that worked best from a preliminary screen (Fig. 4A, lane 5; Supplemental Fig. 6), we observed that the trio combination had the strongest additive effect on exon-16^A333G^ splicing (Fig. 4A, lane 6). Quantitatively, relative to the exon-16^A333G^ control with no ASOs (PSI = 19.1%), the trio ASO combination of ASO_55-72_, ASO_73-90_ and ASO_91-108_ enhanced splicing of exon-16^A333G^ the most significantly (PSI = 77.8%, *P*-value <0.0001), increasing the inclusion of exon-16 by ∼4.1-fold (Fig. 4B). The trio ASO combination completely restores splicing of exon-16^A333G^ to WT levels.

**Figure 4.**
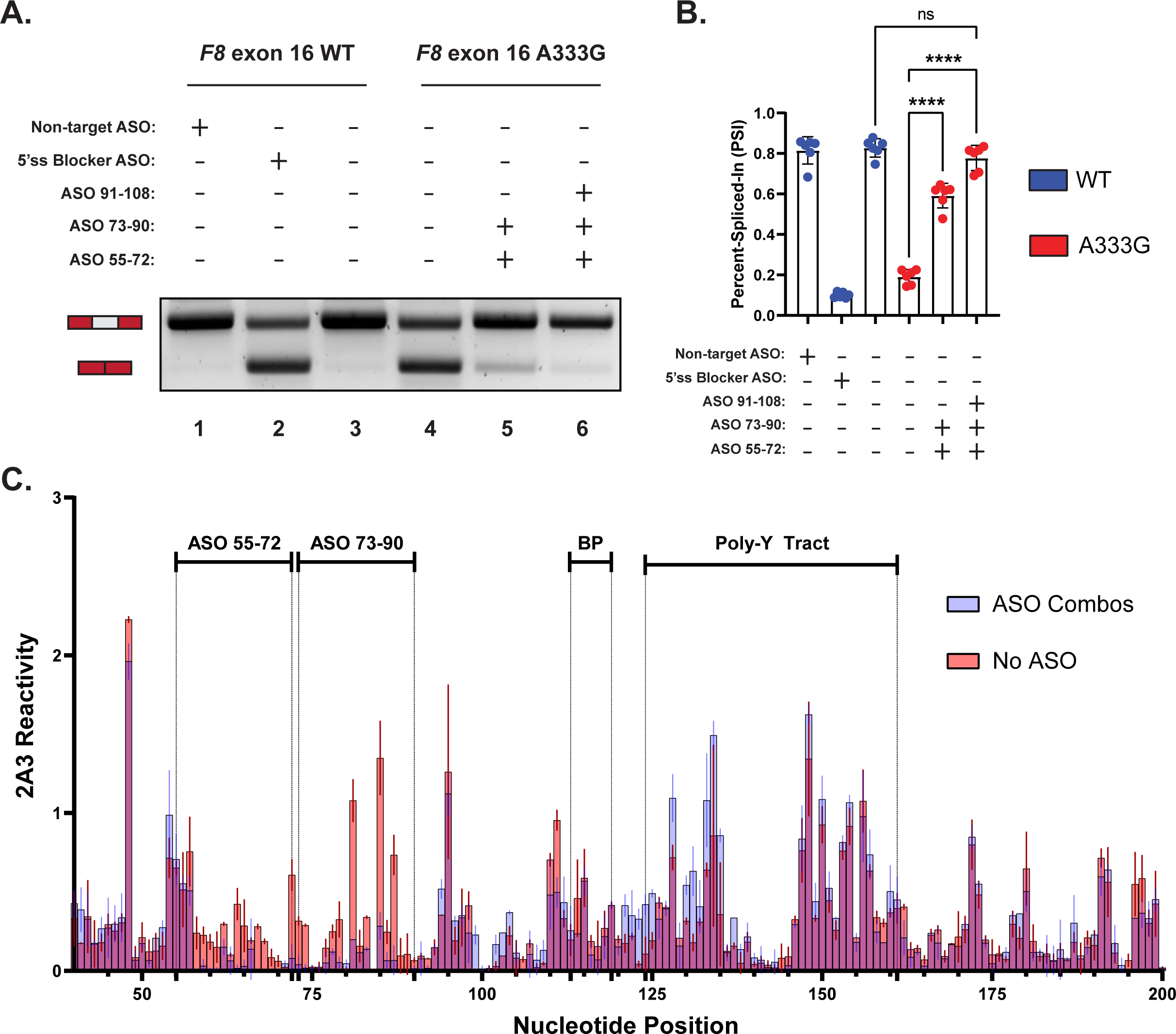
A combination of ASOs targeting TWJ-3-15 can additively enhance splicing of a highly splicing-sensitive mutation by increasing 3′ss accessibility. (**A**) A representative agarose gel depicting the results from our *in vivo* splicing assays testing duo and trio ASO combinations’ ability to modulate splicing. Each splicing assay condition is annotated as shown in the matrix above the gel. Expected mRNA isoforms including or excluding the test exon are also annotated to the left of the agarose gel. (**B**) A plot quantifying the results from (A) using the PSI ratio. The WT context is annotated by a blue color whereas the exon-16^A333G^ mutant is annotated by a red color. The same annotative matrix seen in (A) is used under the plot to label each ASO condition tested for each context. Statistical significance between comparisons are denoted by asterisks that represent *P*-values with the following range of significance: ns = *P* > 0.05, * = *P* ≤ 0.01, *** = *P* ≤ 0.001, **** = *P* ≤ 0.00001. (**C**) An overlay plot comparing normalized 2A3 reactivities between two distinct SHAPE probing conditions used to probe the exon-16^A333G^ mutant. One SHAPE condition probes exon-16^A333G^ with ASOs present (annotated light blue), and the other condition probes exon-16^A333G^ without ASOs present (annotated light red). Admixing of colors (indicated by purple hue) where this is indistinguishable overlap represents similar SHAPE reactivity values between the two probing conditions at that nucleotide position. The nucleotide positions where the ASOs bind, in addition to important splicing signals, are annotated in the plot. All SHAPE probing data presented were generated *in vitro* using the SHAPE reagent 2A3, and all subsequent data analysis was performed in RNA Framework.

To test the hypothesis that the trio ASO combinations targeting TWJ-3-15 increase the accessibility of the poly-Y tract by antagonizing the TWJ-3-15 structure, we performed additional chemical probing experiments. Briefly, we adapted the same SHAPE-MaP-seq approach as previously described with the exception that prior to SHAPE probing, the *in vitro* RNA template was unfolded and then re-folded in the presence of ASOs. In order to draw comparisons between the probing condition with ASOs present and the control condition without ASOs, we calculated the average SHAPE reactivity for each nucleotide from the respective datasets and plotted them together (Fig. 4C). Differences between each dataset’s SHAPE reactivities are represented by their distinct color annotation, whereas predominant admixing of colors represent minimal differences. Accordingly, the results from this experiment show that there is an increased shift in SHAPE reactivities for nucleotides that surround or comprise the poly-Y tract in the condition with ASO_55-72_ and ASO_73-90_ present, compared to the minus ASO condition (Fig. 4C). The reduced SHAPE reactivities in the region of with ASO_55-72_ and ASO_73-90_ complementarity provide a direct measure of ASO binding to the intended target sites (Fig. 4C). Additionally, a single ASO such as ASO_91-108_ is also capable of increasing the SHAPE activities for nucleotides comprising the poly-Y tract (Supplemental Fig. 7). These results indicate that appropriately designed ASOs enhance exon-16 splicing in part by destabilizing TWJ-3-15 to increase the accessibility of the 3′ss, likely increasing the accessibility of the poly-Y tract to U2AF.

### TWJ-3-15 cooperates with hnRNPA1 to weaken F8 exon-16 definition

Results of our RNA structure probing experiments in the presence and absence of specific ASO combinations suggest that TWJ-3-15 reduces the strength of *F8* exon-16 definition by occluding the poly-Y tract. To determine if TWJ-3-15 may also contain any functional binding sites for RNA-binding proteins (RBPs), we used RBPmap to identify RBP consensus motifs within the structure (40). We found two binding sites for the splicing repressor hnRNPA1 within TWJ-3-15 (Fig. 5A, indicated in red). The first potential binding site is found at nucleotide positions 84-90 (U<u>UAGG</u>GA), and the second motif is found at nucleotide positions 99-105 (C<u>UAAGG</u>A). We term these predicted hnRNPA1 binding sites as ISS-15-1 and ISS-15-2, respectively. Based on published research, these predicted binding sites harbor motifs either identical or highly similar to the hnRNPA1 consensus motif, “UAGG”(43, 44). These predicted hnRNPA1 binding sites are positioned within the three-way junction of TWJ-3-15. Intriguingly, ASO_73-90_ and ASO_91-108_ directly bind ISS-15-1 and ISS-15-2, respectively. These ASOs, when used individually or in combination, antagonized the aberrant splicing of the exon-16^A333G^ variant (Fig. 2D). We hypothesize that there is a structure-function relationship where TWJ-3-15 weakens the accessibility of the 3′ss to U2AF due to structural constraints, in addition to recruiting a pair of hnRNPA1 proteins that cooperates with this structure to amplify inhibitory effects at the 3′ss.

**Figure 5.**
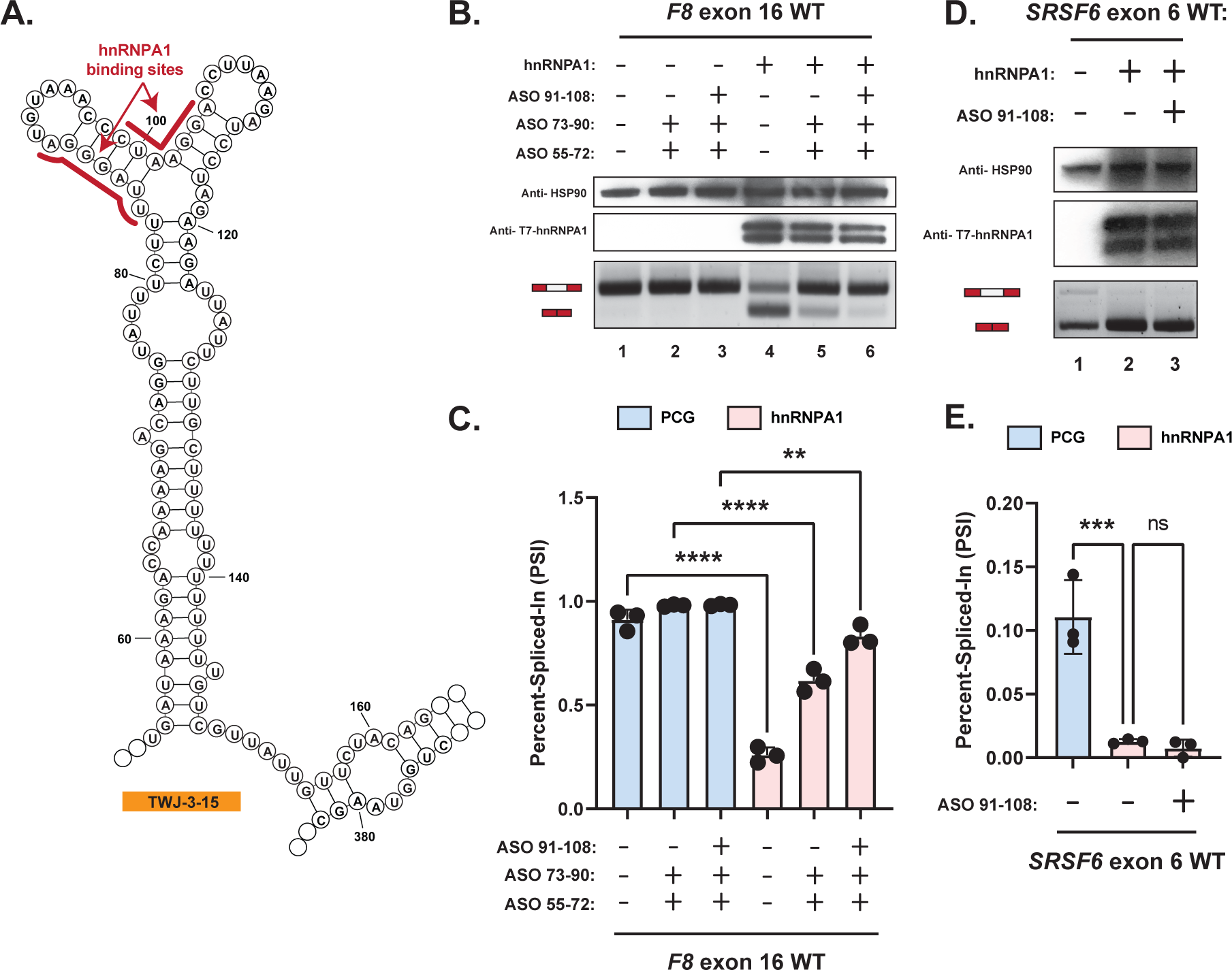
hnRNPA1 cooperates with TWJ-3-15 to amplify inhibitory effects at the 3′ss of *F8* exon-16. **(A)** Secondary structure model of TWJ-3-15 showing predicted hnRNPA1 binding motifs underscored in red. (**B**) Representative Western blot and agarose gel depicting results from our hnRNPA1-ASO competition assay. Each condition tested in the assay is annotated as shown in the matrix above the gel. (**C**) A plot quantifying the results from (A) using the PSI ratio. Co-transfection of the WT exon-16 splicing reporter with either the empty expression vector (PCG) or the hnRNPA1 expression vector is indicated by a light blue or light red color, respectively. The same annotative matrix seen in (A) is used under the plot to label each ASO condition tested for each context. (**D**) Representative Western blot and agarose gel electrophoresis depicting results from our *SRSF6* splicing assay. Each condition tested in the assay is annotated as shown in the matrix above the gel. (**E**) A plot quantifying the results from (C) using the PSI ratio. Co-transfection of the *SRSF6* exon 6 splicing reporter with either the empty expression vector (PCG) or the hnRNPA1 expression vector is indicated by a light blue or light red color, respectively. The same annotative matrix seen in (C) is used under the plot to label each ASO condition tested for each context. Epitopes targeted by specific antibodies in the Western blots are indicated to the left of their respective blots. Expected mRNA isoforms including or excluding the test exon are also annotated to the left of the agarose gel. Statistical significance between comparisons are denoted by asterisks that represent *P*-values with the following range of significance: ns = *P* > 0.05, **\*** = *P* ≤ 0.05, **\*\*** = *P* ≤ 0.01, **\*\*\*** = *P* ≤ 0.001, **\*\*\*\*** = *P* ≤ 0.0001.

To test the hypothesis that promising ASO combinations act to enhance splicing by also blocking hnRNPA1 binding to ISS-15-1 and ISS-15-2 within TWJ-3-15, we performed hnRNPA1-ASO competition assays. ASOs and a splicing reporter were co-transfected with either an empty expression vector or with an hnRNPA1 expression vector. Conditions with the hnRNPA1 expression vector should lead to the overexpression of hnRNPA1, which would be expected to inhibit splicing of exon-16 if the putative hnRNPA1 binding sites are functional. If splicing inhibition directed by hnRNPA1 is attenuated with ASOs that target the predicted silencers ISS-15-1 and ISS-15-2, this would lend support to the notion that hnRNPA1 physically interacts with TWJ-3-15. We chose to test whether a duo or trio combination of ASOs would be more effective in attenuating splicing by targeting the silencers in TWJ-3-15. We used ASO_55-72_ in combination with ASOs that target ISS-15-1 and/or ISS-15-2. We elected to focus this experiment on the WT exon-16 reporter construct to test whether ASOs targeting TWJ-3-15 can attenuate hnRNPA1-dependent inhibition of splicing. The WT reporter has the nearly 100% PSI and therefore will provide the most dynamic range to determine if overexpression of hnRNPA1 can induce exon-16 skipping. Importantly, we note that the predicted hnRNPA1 binding sites are found in the same predicted RNA structure observed within all *F8* exon-16 contexts (WT and mutants) in this study; thus results with the WT construct will also validate the possible ASO mechanism for the mutant exon-16 sequences.

The results of our hnRNPA1-ASO competition assay demonstrate that overexpression of hnRNPA1 can strongly inhibit splicing of WT exon-16 (compare lane to lane 1, Fig. 5B,C). In comparison to the condition where WT exon-16 was co-transfected with the hnRNPA1 expression vector but not with any ASO combinations present (lane 4), we observe that both a duo or trio ASO combination can attenuate the inhibitory effects of hnRNPA1 on exon-16 splicing (compare lane 4 to lane 5 and 6). It is interesting to observe that ASO_55-72_ and ASO_73-90_ are sufficient to attenuate hnRNPA1-directed inhibition of splicing (lane 5). In this case, only ISS-15-1 is sterically masked by ASO_73-90_. In contrast, when analyzing the trio ASO combination where ASO_55-72_ relieves the poly-Y tract and both ISS-15-1 and ISS-15-2 are respectively blocked by ASO_73-90_ and ASO_91-108_, we see the strongest rescue effect where splicing is considerably improved in the presence of hnRNPA1 overexpression (lane 6). ASO_91-108_ contains a potential high affinity hnRNPA1 binding site (5′-AGGTCCT<u>TAGG</u>GTTTACA - 3′), inviting the hypothesis that ASO_91-108_ may attenuate exon-16 splicing in our hnRNPA1-ASO competition assay by directly binding to and sponging hnRNPA1. To test this hypothesis, we repeated the hnRNPA1-ASO competition assay using an orthogonal hnRNPA1-responsive splicing reporter that does not bind ASO_91-108_. This splicing reporter contains *SRSF6* exon-6, a reporter previously validated to exhibit splicing suppression upon hnRNPA1 overexpression (45). We observe that, relative to the empty expression vector, overexpression of hnRNPA1 caused skipping of *SRSF6* exon-6, as expected (Fig. 5D, compare lane 1 to lanes 2 and 3; Fig. 5E). The effect of hnRNPA1 overexpression is maintained in the presence of ASO_91-108_ (Fig. 5D, lane 2), suggesting that ASO_91-108_ does not attenuate hnRNPA1-directed inhibition of splicing by binding to the splicing factor itself. Taken together, these experiments support our hypothesis that promising ASO combinations are indeed blocking *bona fide* hnRNPA1-dependent silencers found within TWJ-3-15.

### A trio ASO combination targeting TWJ-3-15 rescues splicing of multiple HA-linked variants of F8 exon-16

Our experiments indicate that the splicing fidelity of *F8* exon-16 appears to be regulated in part by an RNA structure that weakens the 3′ss and recruits hnRNPA1 to this exon-intron junction (Fig. 6A). Because this feature is shared across all exon-16 mutant alleles (Supplemental Figs. 8-14) and is targeted by ASOs capable of rescuing exon-16^A333G^ splicing, we reasoned that targeting TWJ-3-15 might rescue splicing of other exon-16 mutants. To test this hypothesis, we co-transfected the WT, G351A, C348T, C321A, C321T, and C179T exon-16 splicing reporters with the NT ASO or the trio ASO combination. Similar to exon-16^A333G^, we can observe that co-transfecting ASO_55-72_, ASO_73-90_ and ASO_91-108_ together can rescue the splicing of other splicing-sensitive HA-causing mutants of exon-16, including the WT context (Fig. 6B). Relative to the NT ASO condition for WT exon-16 (lane 1), we can observe that the trio ASO combination can strengthen the splicing efficiency of WT exon-16 splicing in a non-significant manner (lane 2). In comparing the NT ASO condition for each exon-16 mutant (lanes 4, 7, 10, 13 and 16) to their condition with the trio ASO combination co-transfected (lanes 5, 8, 11, 14, and 17), we observed that aberrant splicing of these HA-linked exon-16 variants were significantly ameliorated with all three ASOs present (Fig. 6C). Taken together, these data demonstrate that ASO_55-72_, ASO_73-90_ and ASO_91-108_ interfere with the function of TWJ-3-15 and can generally rescue a broad array of splicing-sensitive mutant alleles.

**Figure 6.**
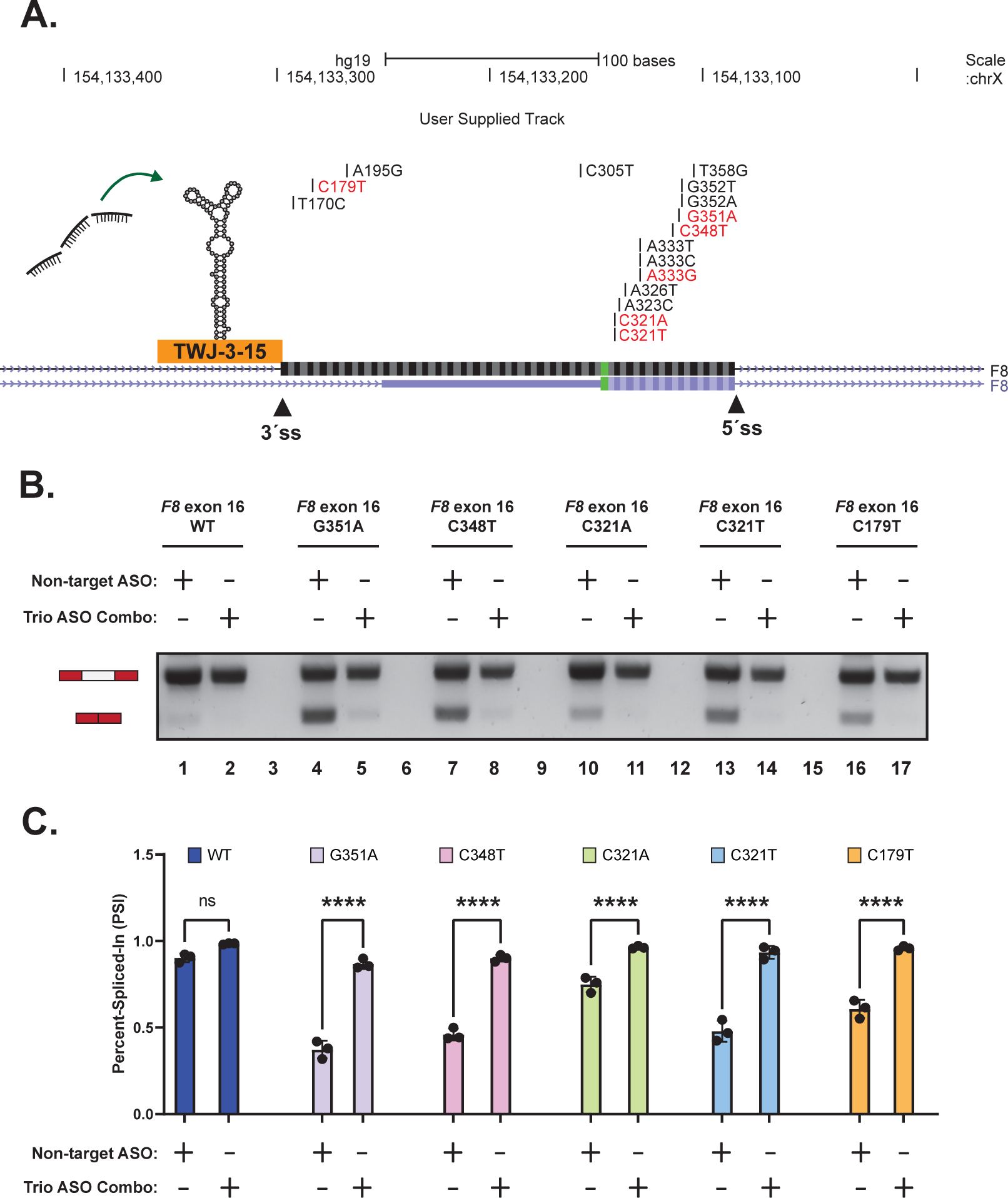
A combination of ASOs targeting TWJ-3-15 can reverse aberrant splicing for a broad array of Hemophilia A associated variants of exon-16 by increasing 3′ss accessibility and blocking hnRNPA1 binding. (**A**) A UCSC Genome Browser screenshot depicting the *F8* exon-16 locus and the positions of HA-causing mutations tested in this study. The 3′ and 5′ splice sites are annotated in addition to TWJ-3-15. Successful ASOs targeting TWJ-3-15 are depicted using the same color scheme as previously shown in Fig. 3B. (**B**) A representative agarose gel depicting the results from our *in vivo* splicing assays testing the trio ASO combinations’ ability to reverse aberrant splicing of exon-16 induced by other HA-causing mutations. Each splicing assay condition included in this specific assay is annotated as shown in the matrix above the gel. Expected mRNA isoforms including or excluding the test exon are also annotated to the left of the agarose gel. (**C**) A plot quantifying the results from (B) using the PSI ratio. Each sequence context tested (WT or MT) is annotated by a distinct color. The same annotative matrix seen in (B) is used under the plot to label each ASO condition tested for each context. Statistical significance between comparisons are denoted by asterisks that represent *P*-values with the following range of significance: ns = *P* > 0.05, **\*** = *P* ≤ 0.05, **\*\*** = *P* ≤ 0.01, **\*\*\*** = *P* ≤ 0.001, **\*\*\*\*** = *P* ≤ 0.0001.

## DISCUSSION

This study explores the dichotomy between “fragile” and “robust” exons. Because fragile exons harbor weak splicing signals, their processing is enhancer-dependent (46). Disease-causing mutations can readily induce aberrant splicing of fragile exons (47). Here we report the discovery of a functional RNA element that may attenuate *F8* exon-16 3’ splice site strength. TWJ-3-15 sequesters the polypyrimidine tract and recruits the splicing repressor hnRNPA1. Despite strong splicing site signals (Supplemental Table 3), TWJ-3-15 sensitizes exon-16 to HA-causing mutations. Using the exon-16^A333G^variant as a model, we discovered ASOs that fully rescue exon-16 inclusion. These splice-switching ASOs target TWJ-3-15 and underscore the functional significance of this sequence. Furthermore, other splicing-sensitive mutations in exon-16 are rescuable by the same ASOs. Finally, targeting TWJ-3-15 increases splicing efficiency of WT exon-16 as well. Our work implicates pre-mRNA structure as an important determinant of exon identity.

There is growing evidence that structured features within pre-mRNA can regulate splicing (48). For example, structural elements can influence splice site accessibility and pairing (49, 50). Protein-RNA interactions can also modulate RNA structure and vice versa (51–53). In the example of *F8* exon-16, TWJ-3-15 weakens the 3′ss through a combination of mechanisms. First, the poly-Y tract forms an extended stem with upstream intronic sequences. Second, the structure positions high affinity hnRNPA1 binding motifs in proximity to the exon-16 3′ss. How exonic splicing enhancers activate the 3′ss and attenuate the function of hnRNPA1 is unknown. It is possible that splicing activators such as SRSF1 antagonize the function of hnRNPA1 as described for HIV *TAT* exon-3 (54), Fig. 7). However, it is also possible that splicing activators could promote an alternative RNA conformation that either masks the hnRNPA1 binding sites or increases the accessibility of the 3′ss and branchpoint.

**Figure 7.**
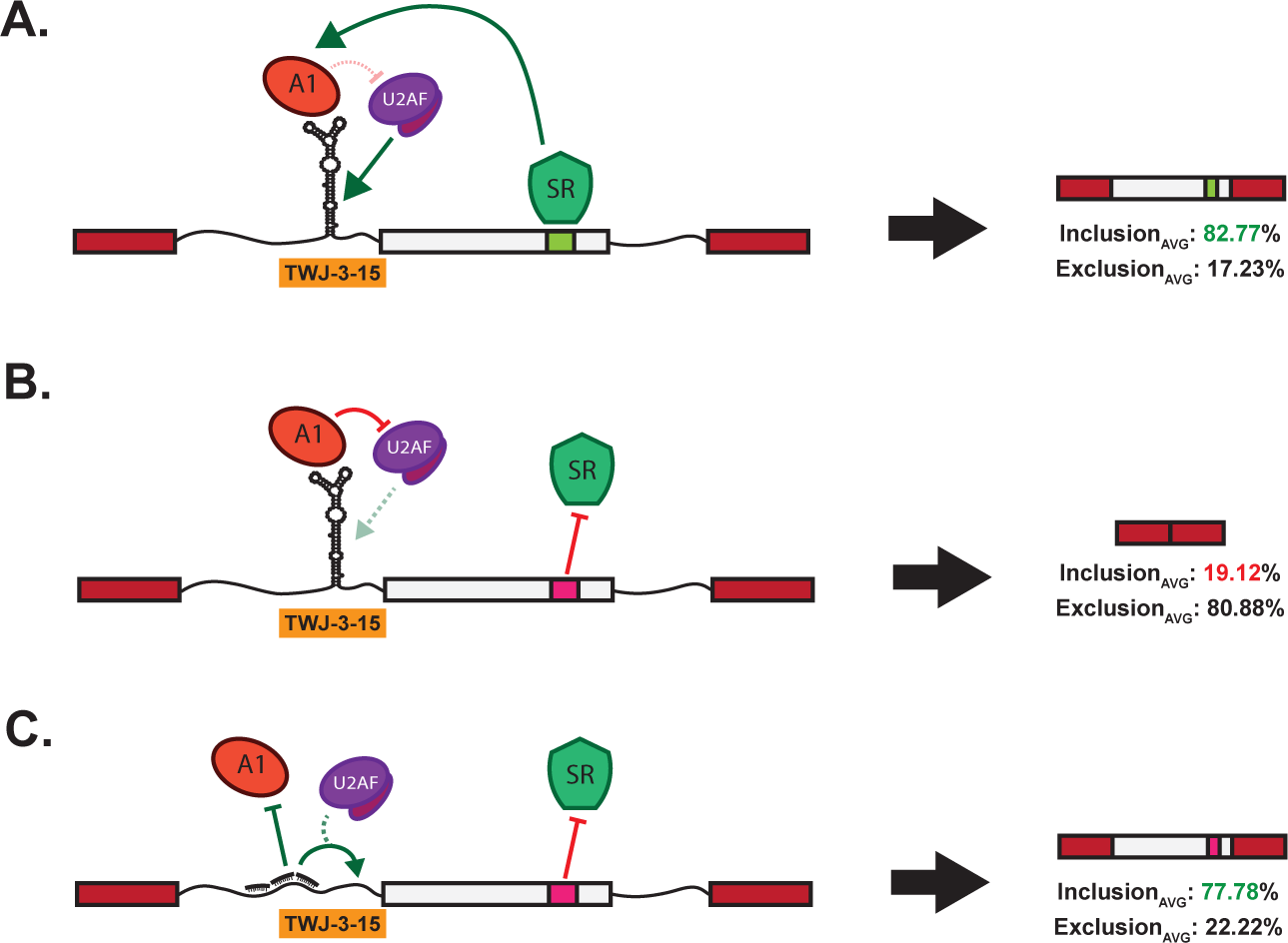
The loss of a critical ESE in *F8* exon-16 presumably amplifies the inhibitory nature of TWJ-3-15 to alter exon definition and splicing fidelity. (**A**) A schematic depicting WT exon-16 (shown in light gray) in our splicing reporter context from in this study. In the WT context, a functional ESE recruits a positive splicing factor that controls the structure-function mechanism comprising TWJ-3-15 and hnRNPA1 at the 3′ss of exon-16. Doing so appears to regulate the inhibitory effects of TWJ-3-15 and hnRNPA1 cooperation, increasing the accessibility of the 3′ss to the splicing machinery. (**B**) A schematic depicting the loss of the ESE in exon-16 due to the A333G mutation. Losing the ESE diminishes the ability to regulate TWJ-3-15 and hnRNPA1 cooperation at the 3′ss exon-16, leading to decreased accessibility of the 3’ss. (**C**) A schematic depicting the trio ASO combinations’ ability to reverse A333G-induced aberrant splicing of exon-16 by destabilizing TWJ-3-15, and preventing the recruitment of hnRNPA1 to the 3′ss. Collectively, our data-supported model indicates that the trio ASOs block the recruitment of a negative splicing factor and to increase the accessibility of the 3’ss to the splicing machinery. TWJ-3-15 is annotated by a simplified depiction of the “Y-shaped” RNA secondary structure at the 3′ss of exon-16. RBPs binding to TWJ-3-15 and this region such as hnRNPA1 and U2AF are respectively annotated. The predicted ESE is annotated in light green within exon-16, and its binding partner, presumably an RBP like SR proteins that are known to enhance splicing, is depicted as well. The loss of the ESE by the A333G mutation is annotated in pink within exon-16.

ASOs are important modalities for precision medicine as they are highly specific and customizable. *F8* exon-16 encodes a portion of the A3 domain which is required for efficient blood clotting and is frequently mutated in HA (55–57). Therefore, TWJ-3-15 is an attractive ASO target in HA. Modulating its function rescues splicing of a diverse array of splicing-sensitive HA-linked variants of exon-16. We also used ASOs to interfere with splicing of the WT *F8* exon-16 reporter. These results demonstrated that all ASOs hybridizing to the exon sequence, spare one, inhibit pre-mRNA splicing. While ASO_199-216_ hybridizes to a sequence predicted to function as a splicing silencer and therefore might not be expected to interfere with splicing, it is less clear why other ASOs targeting the exon are deleterious to splicing. One possibility is that exon-16 is highly structured and that this structure is important for exon definition. Indeed, SHAPE-MaP-Seq suggests extensive intramolecular pairing juxtaposes the 5’ss and 3’ss. It is possible that ASOs targeting exon-16 interfere with base-pairing interactions within the exon and disrupt splicing. Alternatively, *F8* exon-16 may be strongly dependent on exonic splicing enhancers. In this model, ASOs targeting the exon could mask binding sites for splicing factors.

There are several limitations of our study. First, we do not at present understand how HA-causing mutations disrupt exon-16 splicing. One possibility is that point mutations disrupt splicing enhancers (Fig. 7B). Yet, it is possible that mutations remodel RNA structure as suggested by our SHAPE-MaP-seq data. A further limitation of the study is the reliance on splicing reporter assays. Our work utilizes a well-established *HBB* reporter system and HEK293T cells. In this highly-controlled system, exon definition is context dependent as *cis*-regulatory information is often located in proximity to splice sites. However, other factors and processes such as chromatin modification and transcriptional elongation can regulate splicing (58). Testing mutations and ASOs in the context of the endogenous *F8* locus in a disease model is an important next step to reveal if TWJ-3-15 is indeed a *bona fide* target for HA therapies. Finally, it will be important to determine how the *F8* protein tolerates missense mutations in exon-16.

Recent studies demonstrate non-conservative substitutions, including those associated with aberrant splicing, have modest effects on protein activity when expressed from a cDNA (59). Thus, the potential translation of ASOs targeting TWJ-3-15 requires a similar study of *F8* protein function and secretion.

## DATA AVAILABILITY

All sequencing data are available through the Gene Expression Omnibus Short Read Archive (GEO SRA)

## SUPPLEMENTARY DATA

Supplementary Data are available at NAR online.

## AUTHOR CONTRIBUTIONS

V.T., G.C., M.D.S., and J.R.S. conceptualized and led the experiments presented in this study. V.T., H.T., P.H.D., C.O., P.C., I.Q., A.H., and S.L generated the *F8* splicing reporters. V.T. and M.G. performed all cell-based *in vivo* splicing assays. V.T. developed and established the automation platform used in this study. V.T. and M.G. designed and assayed all antisense oligonucleotides targeting *F8* exon-16 using the automation platform. G.C., A.G.J., and V.T. generated the *in vitro* RNA templates for SHAPE-MaP-seq. G.C. and N.M.F. designed, performed, prepared, and analyzed all *in vitro* SHAPE-MaP-seq experiments and sequencing libraries for subsequent RNA structure predictions. V.T. and M.G. performed the hnRNPA1-ASO competition assay to experimentally validate hnRNPA1 binding sites identified in this study. V.T. and G.C. performed all of the data analysis and visualization presented in this study. V.T. wrote the manuscript, with input and review of successive manuscript drafts by G.C., M.D.S., and J.R.S.

## ACKNOWLEDGEMENT

We thank the Toxic RNA Lab (TRL), a curricular undergraduate research experience led by J.R.S., for their help in generating the *F8* splicing reporters. We thank Alexander J. Ritter for writing the Python script that expedited our capacity to quickly design and test antisense oligonucleotides presented in this study. We thank Danny Incarnato for the gift of 2A3 and for their help in getting the RNA Framework analysis pipeline up and running. We thank the Sanford lab, and Gina Mawla for their feedback on manuscript drafts.

---

**Supplemental Figure 1.**
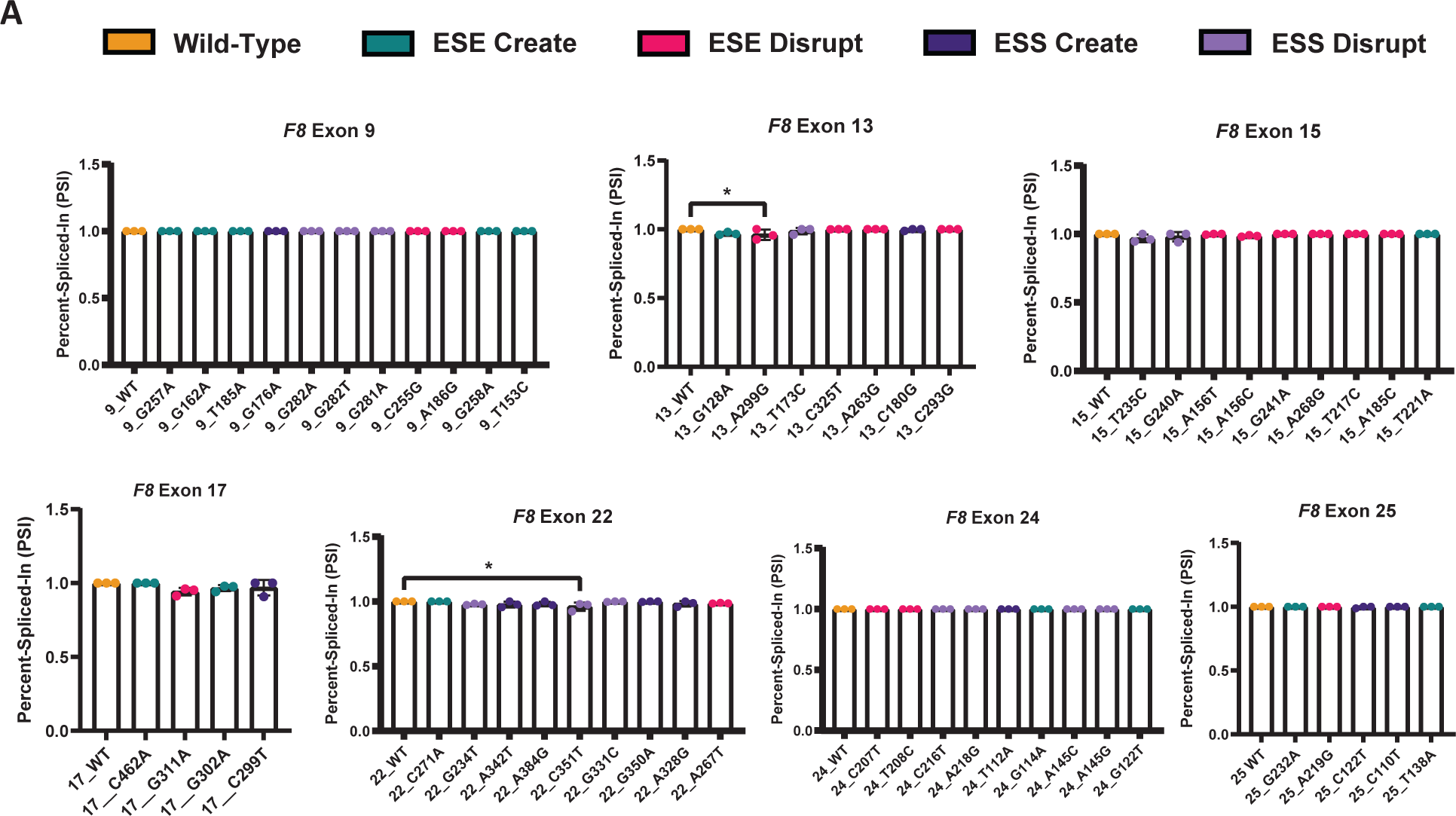
*In vivo* splicing assays reveal robust exons that are resilient to mutation-induced aberrant splicing. **(A)** Percent-spliced-in (PSI) plot quantifying the impact HA-causing mutations have on the splicing of *F8* exons 9, 13, 15, 17, 22, 24, and 25. Each mutation’s predicted impact on regulatory elements involved in splicing are annotated by color. Statistical significance between comparisons are denoted by asterisks that represent P-values with the following range of significance: ns = P > 0.05, * = P < 0.05, ** = P < 0.01, *** = P < 0.001, **** = P < 0.0001.

**Supplemental Figure 2.**
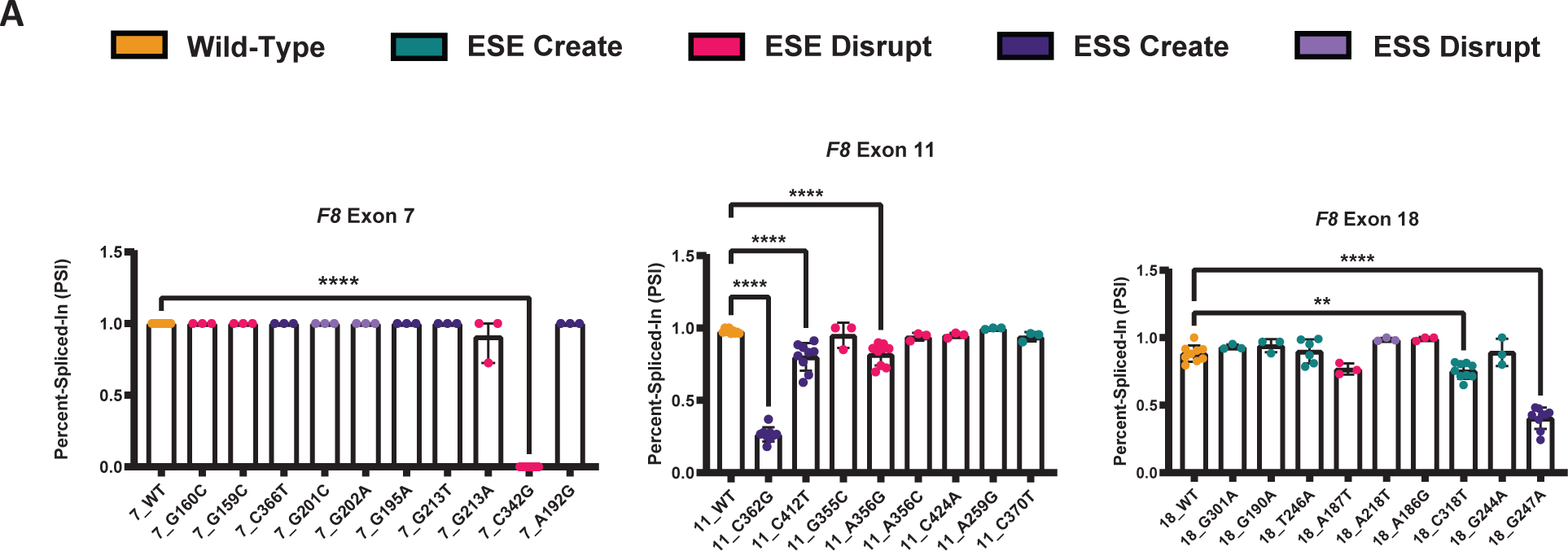
*In vivo* splicing assays reveal fragile exons that are susceptible to mutation-induced aberrant splicing. **(A)** Percent-spliced-in (PSI) plot quantifying the impact HA-causing mutations have on the splicing of *F8* exons 7, 11, and 18. Each mutation’s predicted impact on regulatory elements involved in splicing are annotated by color. Statistical significance between comparisons are denoted by asterisks that represent P-values with the following range of significance: ns = P > 0.05, * = P < 0.05, ** = P < 0.01, *** = P < 0.001, **** = P < 0.0001.

**Supplemental Figure 3.**
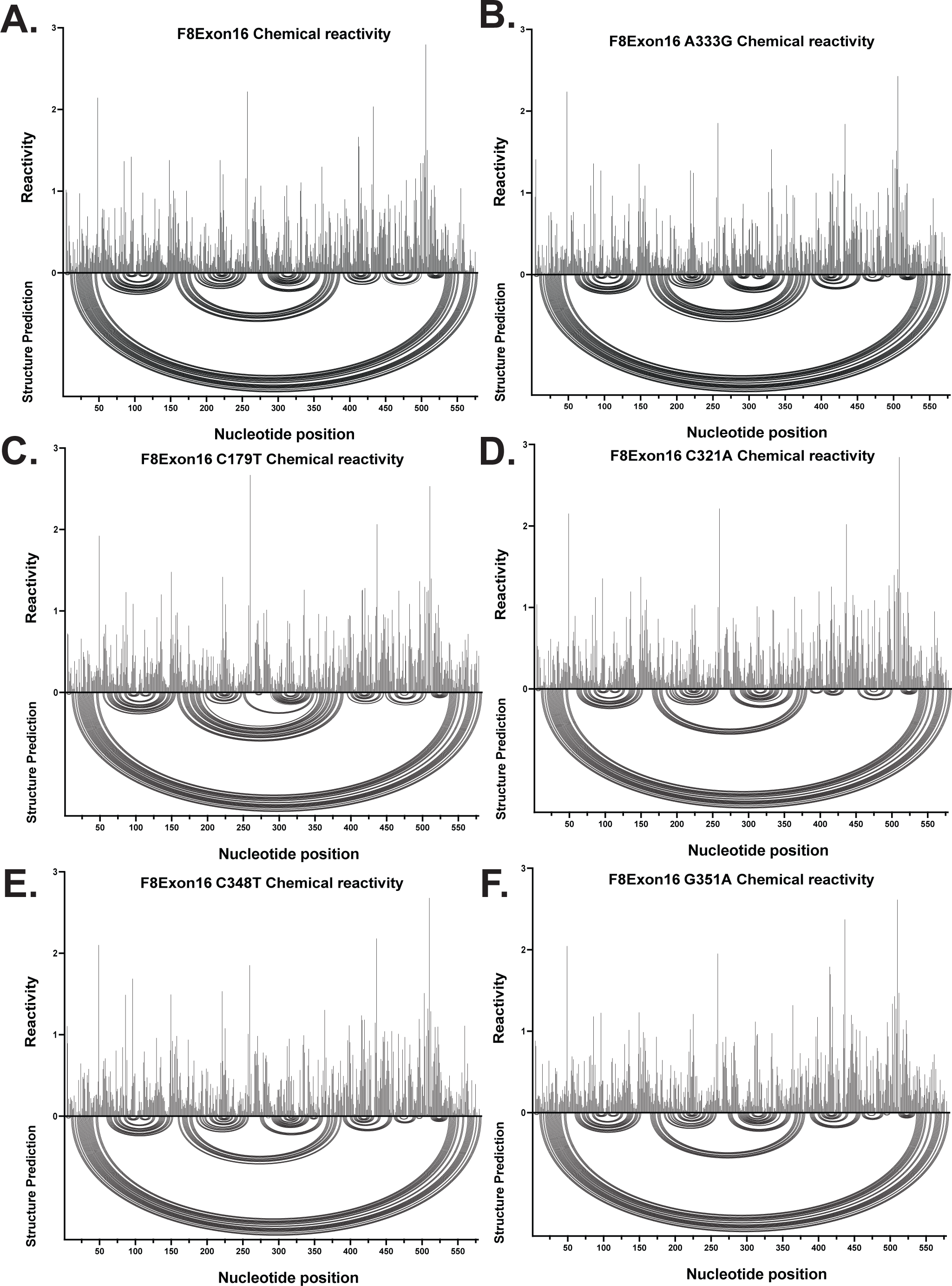
SHAPE reveals that various disease-causing mutations in *F8* exon-16 primarily induce changes to RNA structures proximal to their position. Normalized SHAPE reactivity vs. arc diagram plot comparing WT exon-16 (A) and a HA-causing mutant in their respective panels (B-F). Bars correspond to the normalized SHAPE reactivity for each nucleotide position. The bottom portion of the plot depicts RNA structure predictions using the normalized SHAPE reactivity as a folding constraint. Each arc diagram represents a base-pairing interaction between the respective nucleotides involved within the sequence to form a given RNA structure. **(A)** SHAPE-driven secondary structure prediction for WT *F8* exon-16. **(B)** SHAPE-driven secondary structure prediction for the A333G mutant of *F8* exon-16. **(C)** SHAPE-driven secondary structure prediction for the C179T mutant of *F8* exon-16. **(D)** SHAPE-driven secondary structure prediction of C321A mutant of *F8* exon-16. **(E)** SHAPE-driven secondary structure prediction of C348T mutant of *F8* exon-16. **(F)** SHAPE-driven secondary structure prediction of G351A mutant of *F8* exon-16.

**Supplemental Figure 4.**
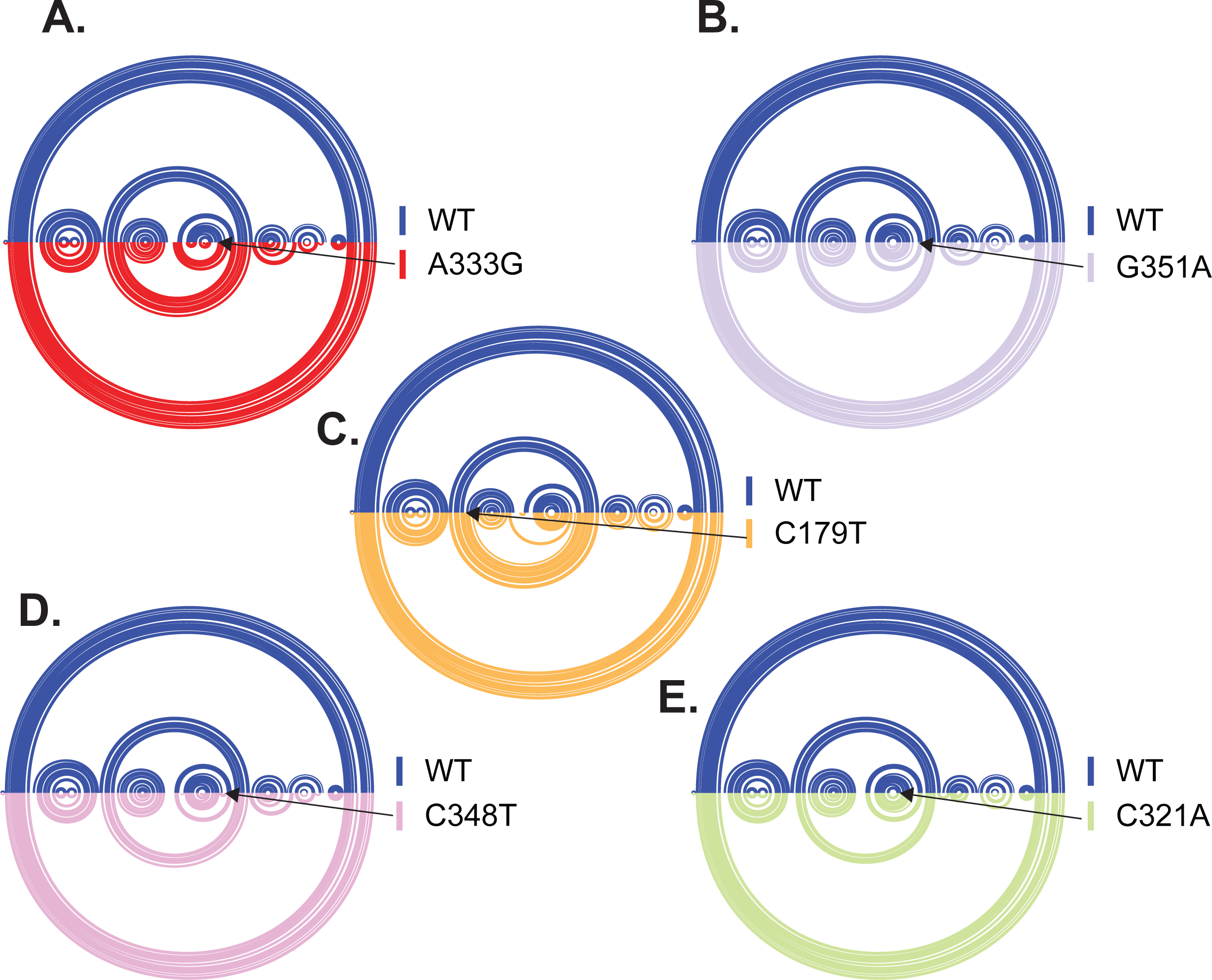
Arc diagrams directly comparing the WT *F8* exon-16 structure prediction to splicing-sensitive mutations. **(A)** Arc diagram comparing WT to the A333G mutation in *F8* exon-16 **(B)** Arc diagram comparing WT to the G351A mutation in *F8* exon-16 **(C)** Arc diagram comparing WT to the C179T mutation in *F8* exon-16. **(D)** Arc diagram comparing WT to the C348T mutation in *F8* exon-16. **(E)** Arc diagram comparing WT to the C321A mutation in *F8* exon-16.

**Supplemental Figure 5.**
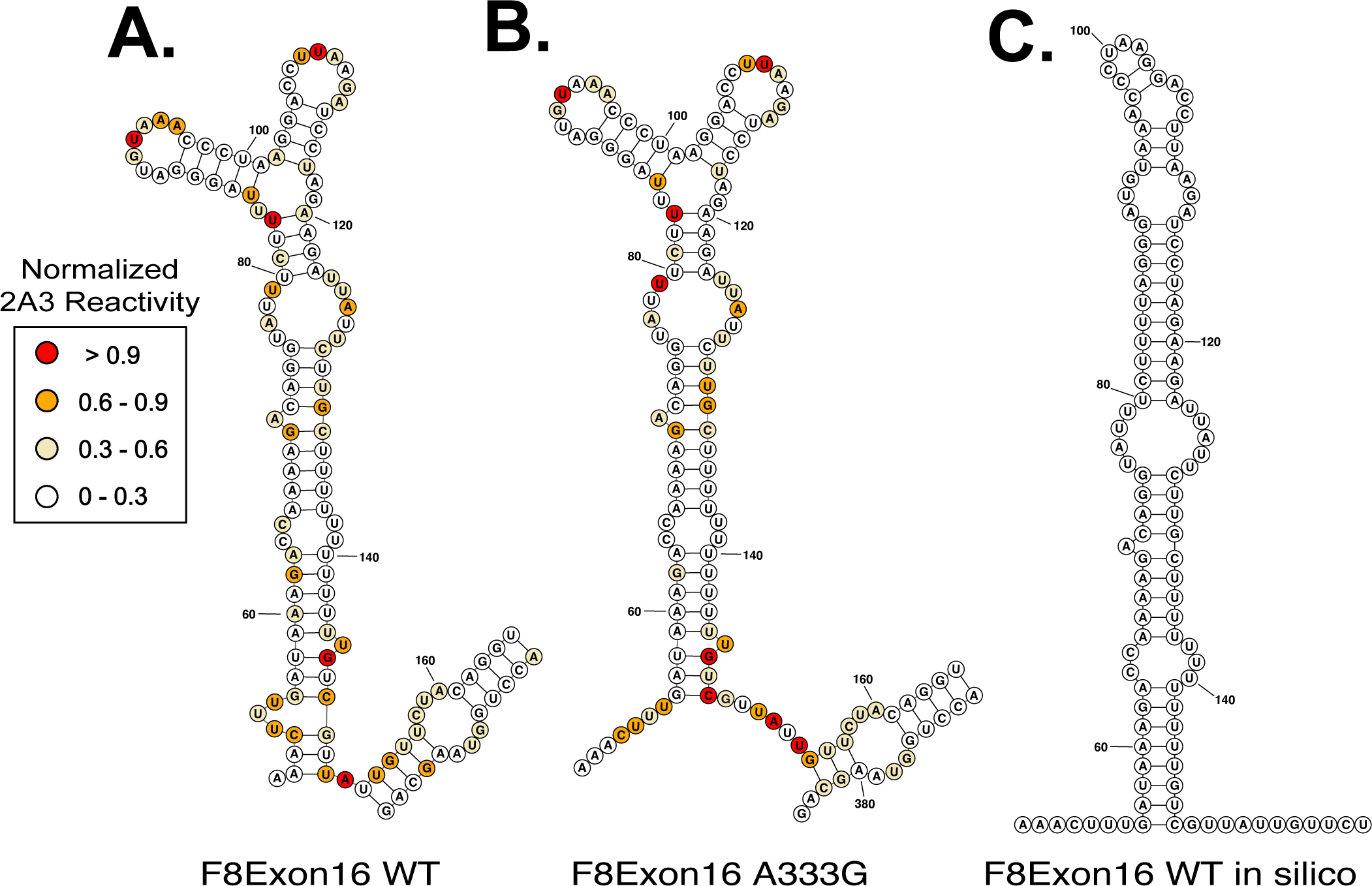
Secondary structure predictions for the sequence corresponding to TWJ-3-15 in wild type or A333G mutant *F8* exon-16 (A and B, respectively). (C) In silico folding of the WT *F8* exon-16 reporter sequence in the absence of SHAPE reactivity data fails to identify TWJ-3-15.

**Supplemental Figure 6.**
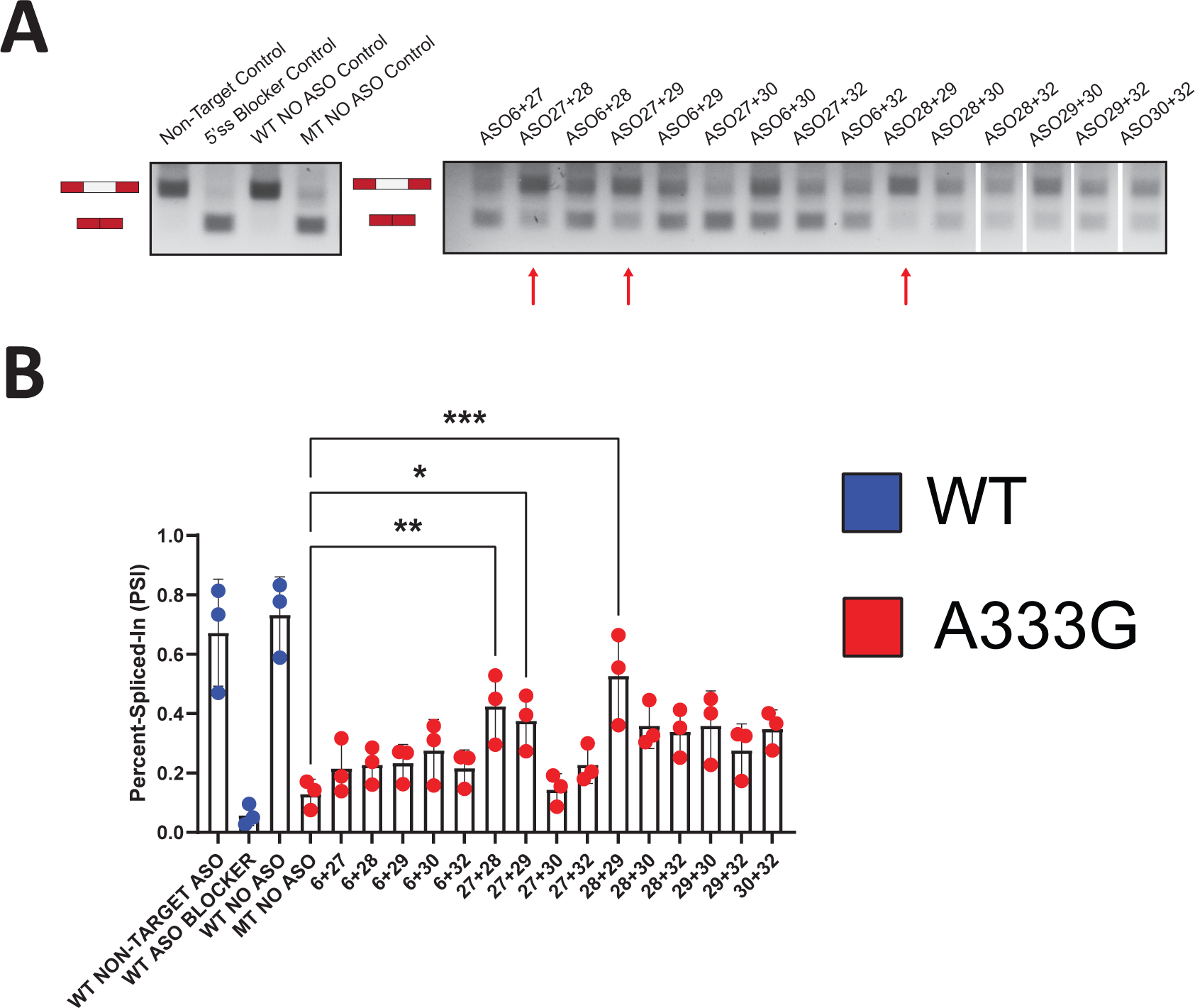
Duo combinations of ASOs targeting *F8* exon-16 can additively enhance splicing of the exon-16^A333G^ mutant (**A**) A representative agarose gel depicting the results from our in vivo splicing assays testing duo ASO combinations’ ability to modulate splicing. Each splicing assay condition is annotated as shown in the matrix above the gel. Expected mRNA isoforms including or excluding the test exon are also annotated to the left of the agarose gel. (**B**) A plot quantifying the results from (A) using the PSI ratio. Each ASO combinations’ ability to significantly modulate splicing is annotated by color and corresponding effect (e.g., enhance or suppress splicing). Statistical significance between comparisons are denoted by asterisks that represent P-values with the following range of significance: ns = P > 0.05, * = P < 0.05, ** = P < 0.01, *** = P < 0.001, **** = P < 0.0001.

**Supplemental Figure 7.**
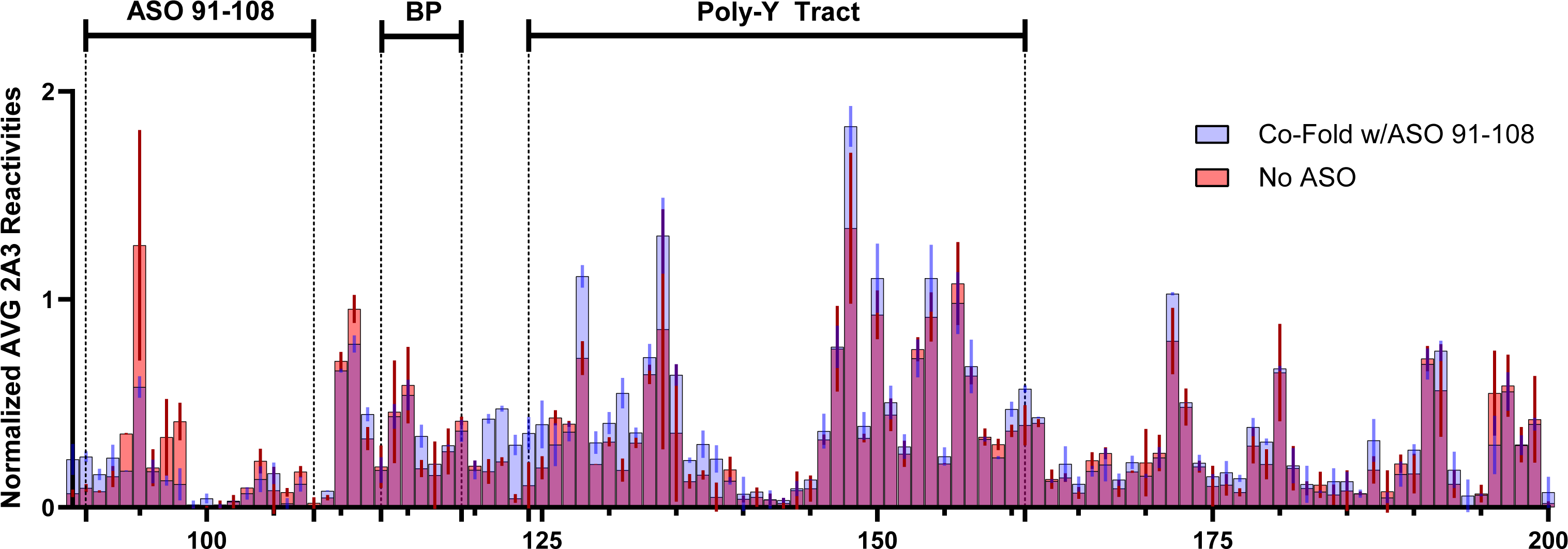
ASOs targeting *F8* exon-16 can increase the accessibility of nucleotides comprising the polypyrimidine tract at the 3’ss. (**A**) An overlay plot comparing normalized 2A3 reactivities between two distinct SHAPE probing conditions used to probe the exon-16^A333G^ mutant. One SHAPE condition probes exon-16^A333G^ with an ASO present (annotated light blue), and the other condition probes exon-16^A333G^ without ASOs present (annotated light red). Admixing of colors (indicated by purple hue) or where there are indistinguishable overlaps represents similar SHAPE reactivity values between the two probing conditions at that nucleotide position. The nucleotide positions where the ASOs bind, in addition to important splicing signals, are annotated in the plot.

**Supplemental Figure 8.**
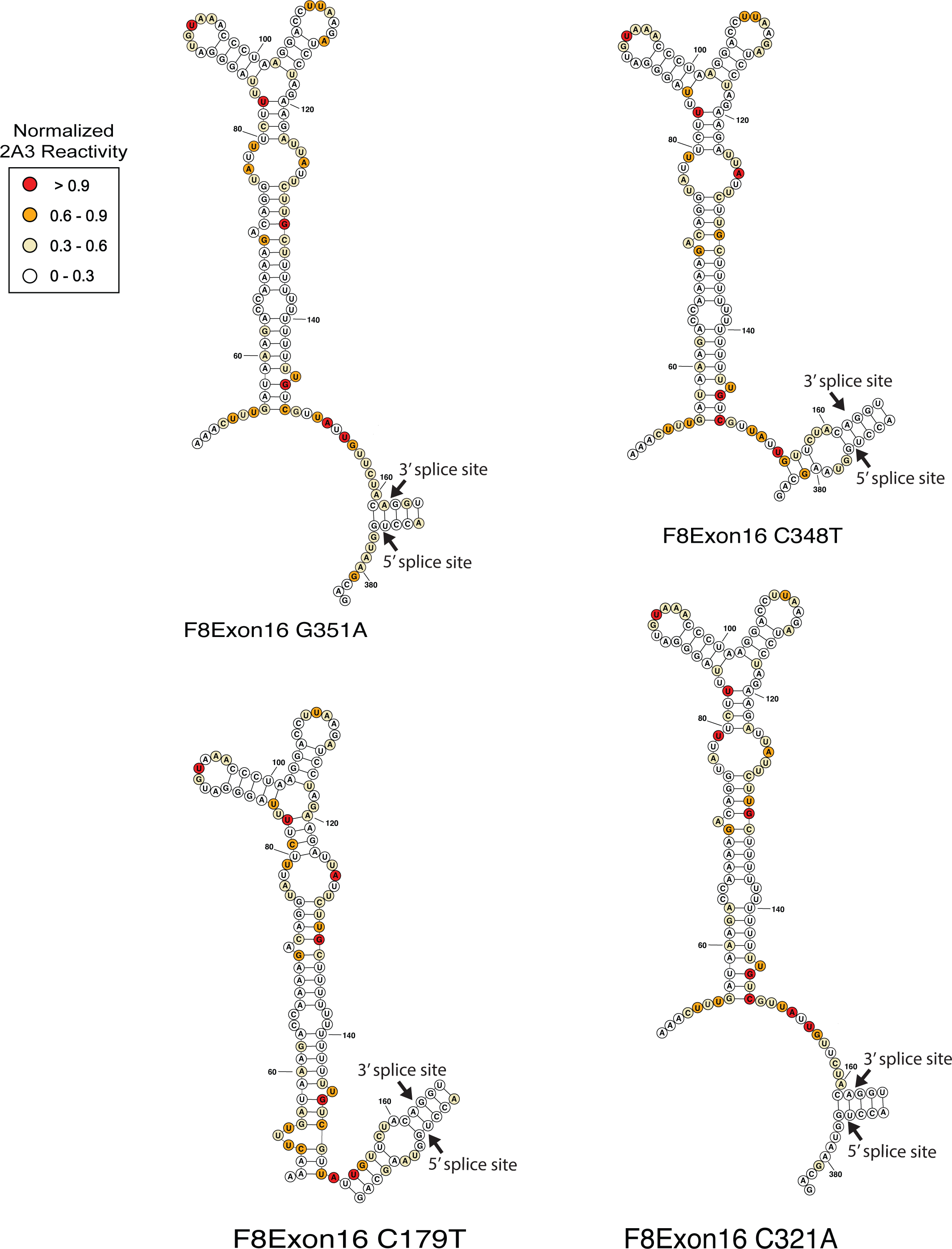
SHAPE-derived secondary structural models for TWJ-3-15 for four additional splicing sensitive alleles. Bases are colored according to their normalized 2A3 reactivity.

**Supplemental Figure 9.**
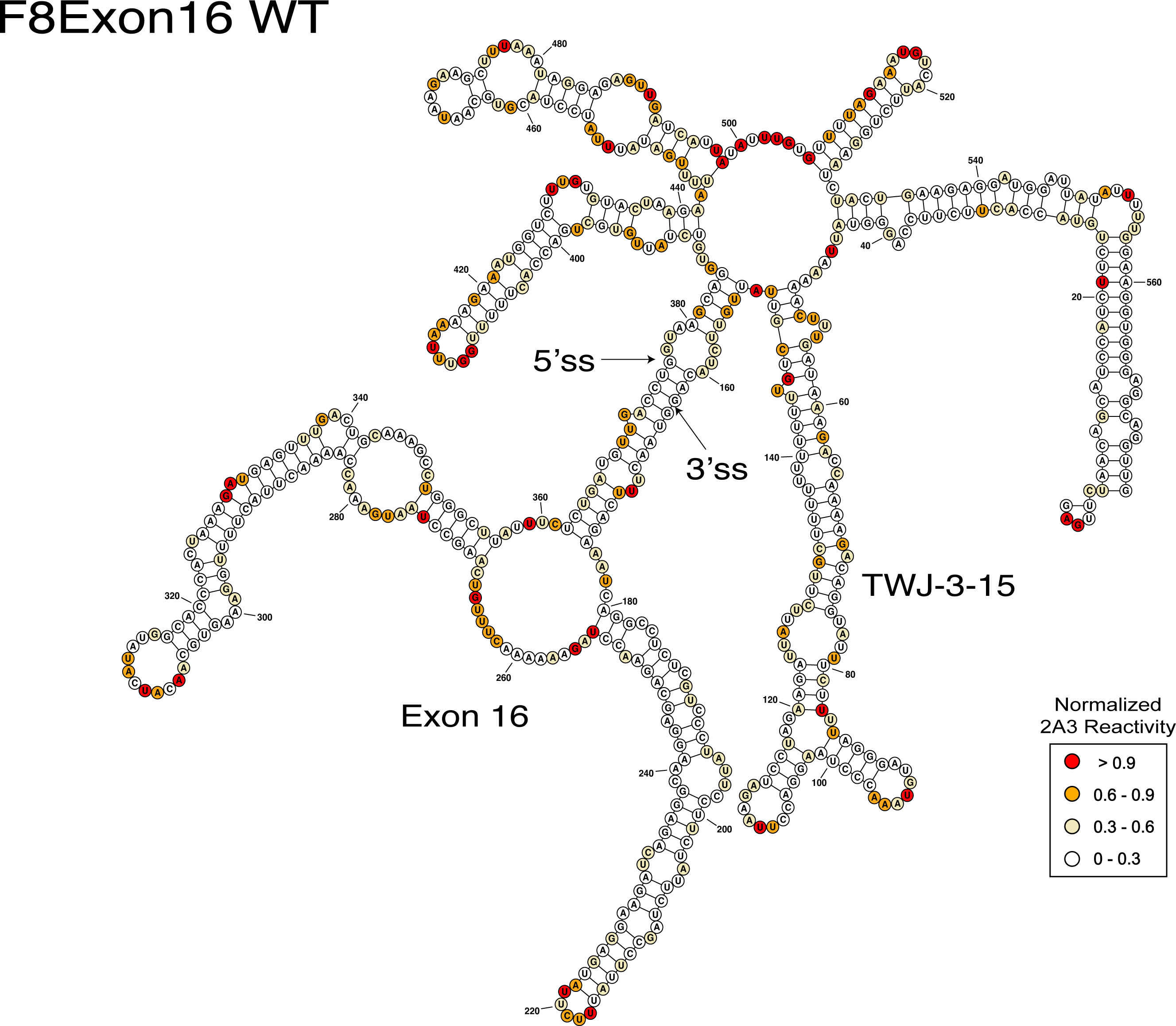
SHAPE-derived secondary structural models for wild type *F8* exon-16 and flanking intron sequences used in splicing reporter assays. Bases are colored according to their normalized 2A3 reactivity.

**Supplemental Figure 10.**
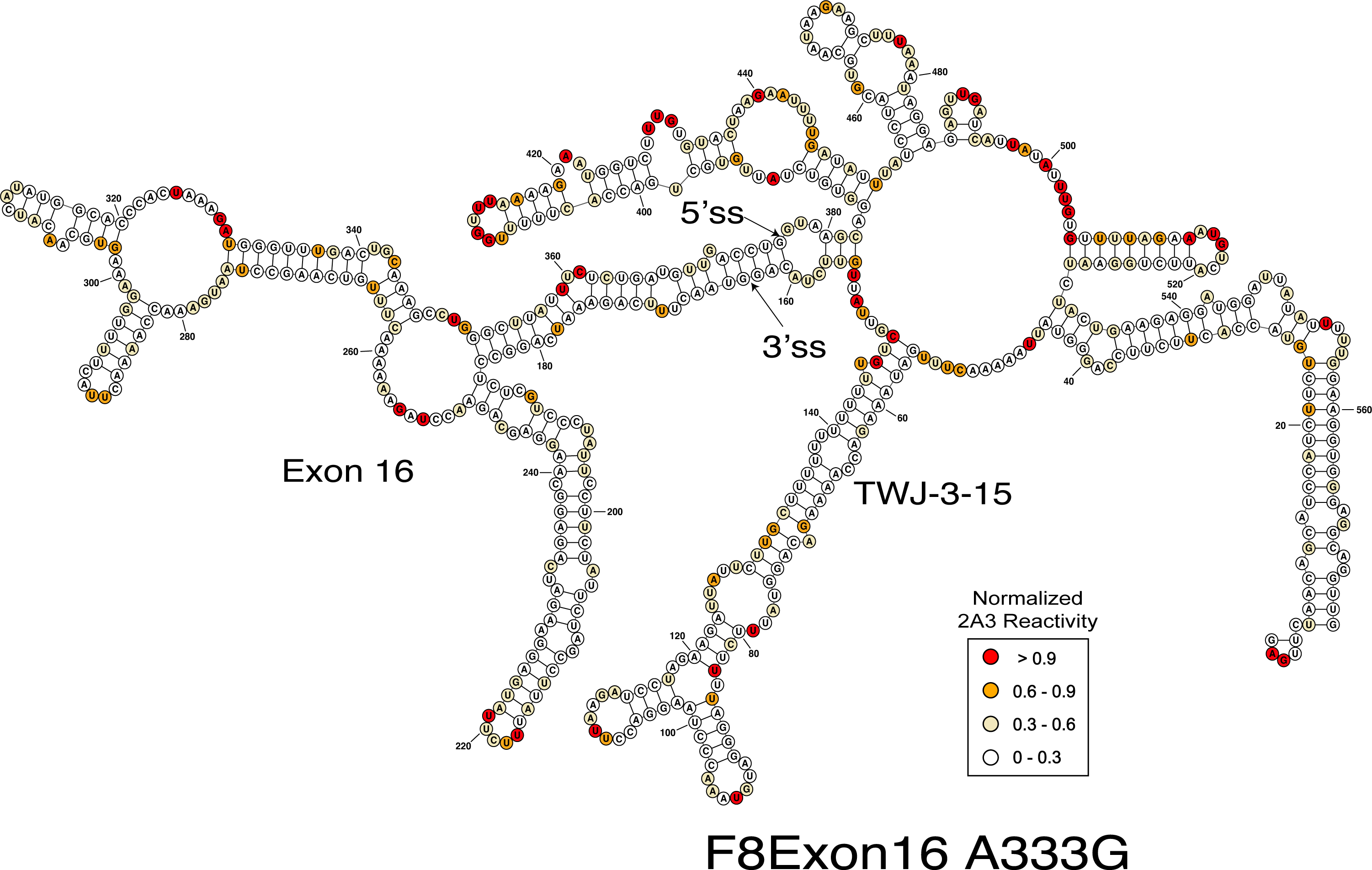
SHAPE-derived secondary structural models for *F8* exon-16^A333G^ and flanking intron sequences used in splicing reporter assays. Bases are colored according to their normalized 2A3 reactivity.

**Supplemental Figure 11.**
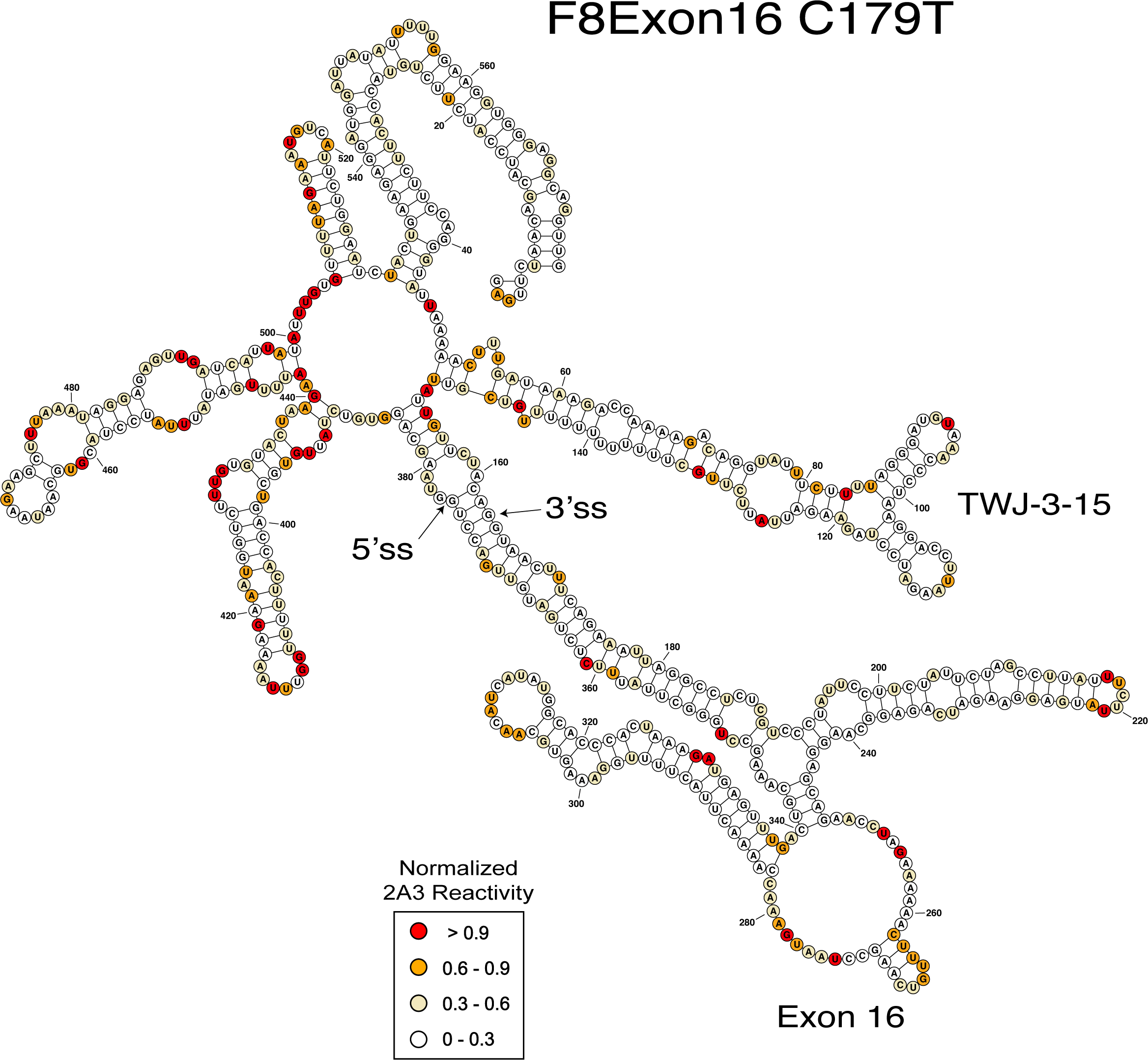
SHAPE-derived secondary structural models for *F8* exon-16^C179T^and flanking intron sequences used in splicing reporter assays. Bases are colored according to their normalized 2A3 reactivity.

**Supplemental Figure 12.**
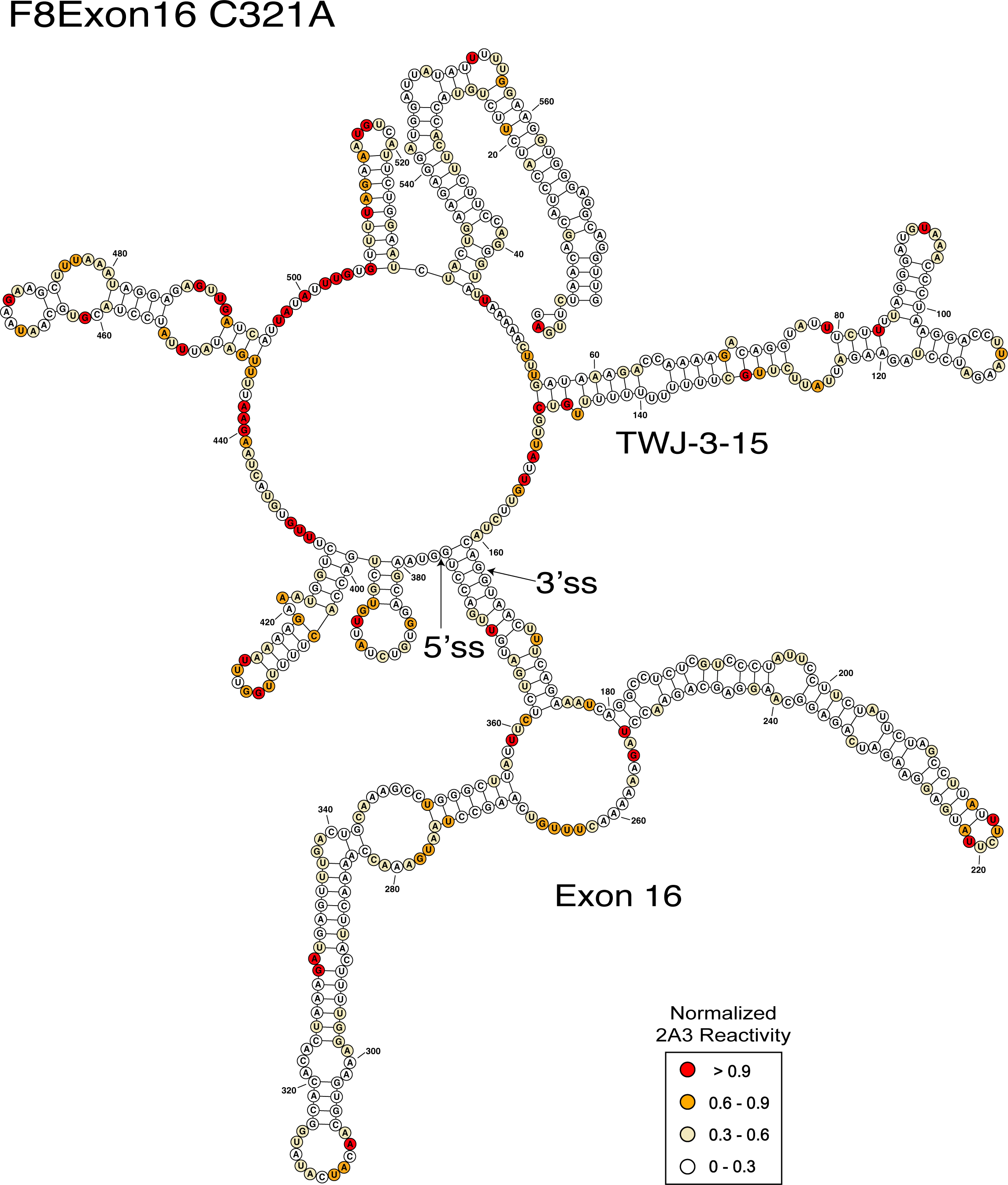
SHAPE-derived secondary structural models for *F8* exon-16^C321A^ and flanking intron sequences used in splicing reporter assays. Bases are colored according to their normalized 2A3 reactivity.

**Supplemental Figure 13.**
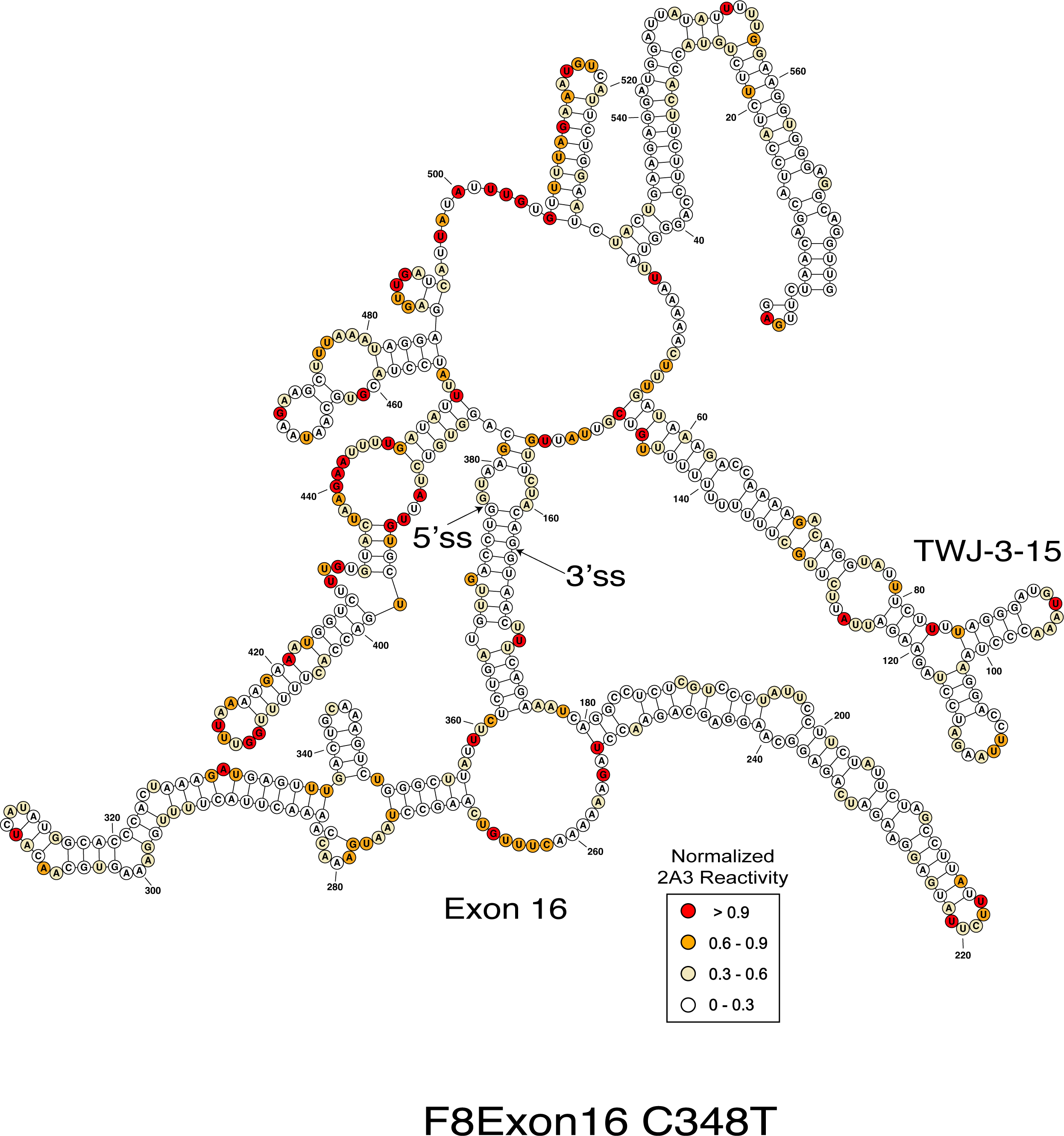
SHAPE-derived secondary structural models for *F8* exon-16^C348T^ and flanking intron sequences used in splicing reporter assays. Bases are colored according to their normalized 2A3 reactivity.

**Supplemental Figure 14.**
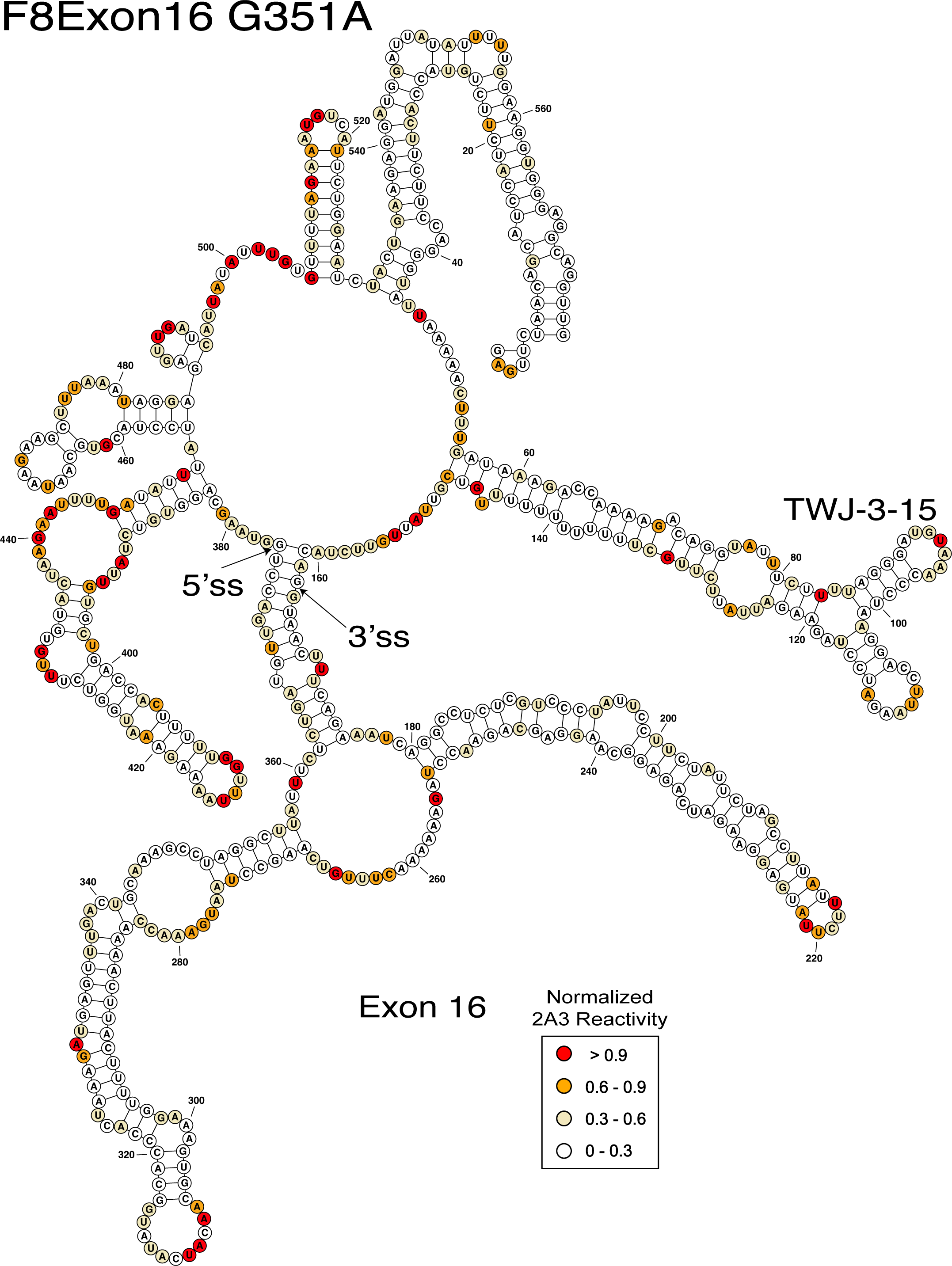
SHAPE-derived secondary structural models for *F8* exon-16^G351A^ and flanking intron sequences used in splicing reporter assays. Bases are colored according to their normalized 2A3 reactivity.

